# A synthetic chlorophagy receptor promotes plant fitness by mediating chloroplast microautophagy

**DOI:** 10.1101/2025.02.21.639444

**Authors:** Rui Liu, Xun Weng, Xinjing Li, Yongheng Cao, Qiyun Li, Lin Luan, Danqing Tong, Zhaosheng Kong, Hao Wang, Taotao Wang, Qingqiu Gong

**Affiliations:** State Key Laboratory of Microbial Metabolism & Joint International Research Laboratory of Metabolic and Developmental Sciences, School of Life Sciences and Biotechnology, Shanghai Jiao Tong University, Shanghai, 200240, China, P. R; Guangdong Provincial Key Laboratory for the Developmental Biology and Environmental Adaption of Agricultural Organisms, College of Life Sciences, South China Agricultural University, Guangzhou, China, P. R; State Key Laboratory of Plant Genomics, Institute of Microbiology, Chinese Academy of Sciences & University of Chinese Academy of Sciences, Beijing 100101, China, P. R; Department of Plant Biology and Ecology, College of Life Sciences, Nankai University, 94 Weijin Road, Tianjin 300071, China, P. R

**Keywords:** Autophagy, chloroplast, chloroplast division, receptor, vacuole

## Abstract

Chloroplasts are major photosynthetic and protein-containing organelles in green plants and algae. Unwanted chloroplast proteins and entire chloroplasts are cleared through various degradation pathways including autophagy. Nevertheless, canonical chlorophagy receptors remain unidentified, and whether and to what extent chlorophagy can be enhanced to benefect the plants remain unknown. Here we designed and validated a synthetic chloroplast autophagy receptor using biochemical, genetical, and imaging approaches. This synthetic receptor, LIR-SNT-BFP, was constructed by fusing a fragment containing the LC3-interacting region (LIR) of the selective autophagy receptor NBR1 and the N-terminal amphipathic α-helix of the chloroplast outer envelope protein SFR2. The fusion protein LIR-SNT-BFP coated the chloroplast and attracted ATG8a *in planta*. Upon induction, the synthetic receptor elicited vacuole-mediated microautophagy of entire chloroplasts independent of the ATG8 conjugation machinery proteins ATG5 or ATG7. Meanwhile, it induced chloroplast division; however the induced chlorophagy was independent of PDV2. Notably, moderate chlorophagy improves rosette growth, but excessive levels are detrimental. Furthermore, induced chlorophagy partially protects against herbicide-induced leaf chlorosis. This study demonstrates controlled chloroplast degradation using a synthetic chlorophagy receptor.

## INTRODUCTION

Eukaryotic cells pack unwanted macromolecules and organelles into double-membraned autophagosomes (macroautophagy) or vacuoles (microautophagy) to maintain homeostasis, and to recycle the building blocks for reuse. Comparing with the ubiquitin-26S proteasome pathway and other intracellular degradation pathways that rely on 26S proteasome, autophagy has much larger capacity and less stringent cargo selection (Raffeiner *et al*, 2023). These characteristics make autophagy an ideal platform for the targeted degradation of large objects. In autophagy, the cargoes for degradation are selectively recognized by autophagy receptors/adaptors, such as NBR1 and p62/sequestosome 1 (SQSTM1) in animals, Atg19 in *Saccchomyces cerevisiae*, and NBR1 in Arabidopsis (Gubas & Dikic, 2022; Yan *et al*, 2024). Ubiquitination and protein aggregation are involved in the process of cargo selection. Mechanistically, the LC3-interacting region/ATG8-interacting motif (LIR/AIM) of autophagy receptors, with a consensus [W/F/Y]xx[L/I/V] (x being any amino acid) surrounded by proximal acidic residues, bind two hydrophobic binding pockets in ATG8/LC3s (Noda *et al*, 2010; Rogov *et al*, 2023). Concurrently, proteins encoded by autophagy-related genes (ATGs) tag the membrane precursor of autophagosome, phagophore/isolation membrane, with ATG8/LC3s, so that autophagy receptors can bring cargoes to phagophore (Yamamoto *et al*, 2023).

Accumulating knowledge of selective autophagy has led to the development of targeted autophagic degradation of proteins and organelles, mainly in mammalian cells and animal models. Autophagosome tethering compounds (ATTECs), identified via high-throughput screening, can bind both LC3s and the target protein, such as mutant huntingtin protein (mHTT), to mediate autophagic degradation of the target protein (Li *et al*, 2019). Likewise, a compound mT1 binds both the mitochondrial outer membrane protein TSPO and LC3B, thus facilitating the autophagic degradation of damaged mitochondria (Tan *et al*, 2023). The autophagy-targeting chimeras (AUTACs) consist of a guanine tag and a specific binder, i.e. HaloTag ligand, which binds a target protein fused with a HaloTag. The guanine tag can trigger K63-linked poly-ubiquitination of cargo proteins, which are recognized by cargo receptors for autophagic degradation (Takahashi *et al*, 2019). Another autophagy-targeting chimera (AUTOTAC) are compounds that bind the autophagy receptor p62. AUTOTAC-bound p62 undergoes conformational changes that exposes its PB1 domain and LIR motif, which facilitates its self-polymerization and LC3 binding, respectively, leading to sequestration and degradation of oncoproteins and aggregates in neurodegeneration (Ji *et al*, 2022). Tethering ATG16L1 or LC3 with ATG16L1-binding peptide (ABP) or LIR can induce targeted degradation of mHTT, and fusion proteins containing mitochondria-targeting sequence of the mitochondrial outer membrane protein TOMM20 plus ABP or LIR promoted degradation of damaged mitochondria (Mei *et al*, 2023). Similarly, LIR of Arabidopsis NBR1 can serve as a tool that links targets to ATG8, and this autophagy receptor can facilitate peroxisome degradation in transiently transformed *Nicotiana benthamiana* leaf epidermal cells, and degradation of proteins in Arabidopsis transgenic lines (Luo *et al*, 2023b).

The chloroplast is a plant- and green algae-specific photosynthetic organelle. The large number (up to 100 per mesophyll cell), size (5 to 10 µm long, much larger than mitochondria and peroxisomes), high protein content (RubisCO can account for 50% of total protein) of chloroplasts make them a natural substrate for autophagy, and an ideal target of autophagy manipulation in plants. Many forms of chloroplast macroautophagy and microautophagy have been discovered, with various stimuli and substrates identified (Izumi & Nakamura, 2018; Zhuang & Jiang, 2019). Several forms require ATG5 and ATG7, the components of ATG8 conjugation machineries. These include vacuolar degradation of RubisCO-containing body (RCB), which contains stroma and envelope proteins but not chlorophyll, and triggered by carbon starvation and leaf senescence (Ishida *et al*, 2008; Izumi *et al*, 2024); starch granule-like structure, which contributes to leaf starch degradation (Wang *et al*, 2013); and the ATG8-INTERACTING PROTEIN1 (ATI1)-labelled plastid body (ATI-PS body), which contains thylakoid, stroma, and envelope proteins and is stimulated by leaf senescence, carbon starvation, and salt stress (Michaeli *et al*, 2014). Apart from the autophagic degradation of chloroplast fragments or components, damaged entire chloroplasts can be degraded by autophagy, either dependent or independent of ATG5 or ATG7. Entire chloroplasts from individually darkened leaves (IDLs) or photo-damaged entire chloroplasts can be transported to the vacuole for degradation, dependent on ATG5 and ATG7 (Izumi *et al*, 2017; Nakamura *et al*, 2018; Ono *et al*, 2013; Wada *et al*, 2009). In contrast, singlet oxygen-induced chloroplast protrusion into the vacuole, observed in the *plastid ferrochelatase two (fc2)* mutant, does not require ATG5 or ATG7 (Lemke *et al*, 2021). NBR1 can be recruited to the surface and interior of photo-damaged chloroplasts covered with ubiquitin, and NBR1-decorated chloroplasts are engulfed by the vacuole in a microautophagy-type process, *i.e.*, independent of ATG7 (Kikuchi *et al*, 2020; Lee *et al*, 2023). In addition to autophagy, senescence-associated vacuoles (SAVs), which contain stroma and triggered by leaf senescence (Carrión *et al*, 2013; Martinez *et al*, 2008; Otegui *et al*, 2005), and chloroplast vesiculation-containing vesicles (CCVs), which contain stroma, thylakoid, and envelope proteins, and triggered by senescence as well as salt and oxidative stresses (Pan *et al*, 2023; Wang & Blumwald, 2014; Žnidarič *et al*, 2025), are also forms of vesicle-mediated chloroplast degradation. These discoveries underscored the importance and complexity of chloroplast vesicular degradation and autophagy. So far, a canonical chloroplast autophagy receptor, which presumably localizes to the outer envelope of chloroplast and recruits ATG8 upon autophagy induction conditions, remains unidentified. Meanwhile, whether chloroplast autophagy can be enhanced, to which degree it can be enhanced, and whether the enhancement can benefit plant growth or stress tolerance, are still open questions.

Here, we addressed these questions by designing and validating an inducible, synthetic chloroplast autophagy receptor. The effects of the induced expression of the synthetic receptor are presented at the sub-cellular, cellular, and organism level. We found that moderate induction of chloroplast autophagy leads to larger rosette sizes. However, the induced autophagy has a clear upper limit for its benefit, beyond which growth inhibition occured. Additionally, this induced autophagy can partially protect the seedlings from herbicide-induced chlorophyll damage and leaf chlorosis. This form of chloroplast autophagy appears to be a type of microautophagy, as it is independent of the ATG8 conjugation machinery, specifically ATG5 and ATG7, and its association with stricking changes in vacuole dynamics. Interestingly, this induced chloroplast autophagy is accompanied by chloroplast division. However, induced chloroplast autophagy was still observed in the *pdv2* mutant, which is characterized by giant chloroplasts. These observation led us to speculate that a novel form of microautophagy may be responsible for this chloroplast degradation.

## RESULTS

### Design and validation of a synthetic chloroplast autophagy receptor

A functional synthetic chloroplast autophagy receptor is expected to localize to the outer envelope (OE) of the chloroplast and recruit ATG8 once expressed. To achieve this, we cloned a fragment containing the LC3 interacting region/ATG8 interacting motif (LIR/AIM) of Arabidopsis NBR1 (**Fig. S1A**), the well-characterized selective autophagy receptor/adaptor (Svenning *et al*, 2011). Then we cloned the N-terminal amphipathic α-helix (SNT) of SENSITIVE TO FREEZING 2 (SFR2) (**Fig. S1B and C**), which targets to the OE of chloroplast (Fourrier *et al*, 2008), and fused it after LIR. For imaging, a blue fluorescent protein (BFP) tag was fused in frame after SNT (**Fig. 1A**). The fusion protein LIR-SNT-BFP coated the chloroplast and co-localized with ATG8a in transiently transformed *N.benthamiana* leaf epidermal cells (**Fig. 1B**) and in transiently transformed Arabidopsis mesophyll protoplasts (**Fig. 1C**). For inducible expression in transgenic Arabidopsis, a glucocorticoid receptor-based inducible gene expression system (GVG) was fused in frame before the receptor (Aoyama & Chua, 1997), resulting in *GVG:LIR-SNT-BFP*, the inducible synthetic chlorophagy receptor (**Fig. 1A**).

**Figure 1.**
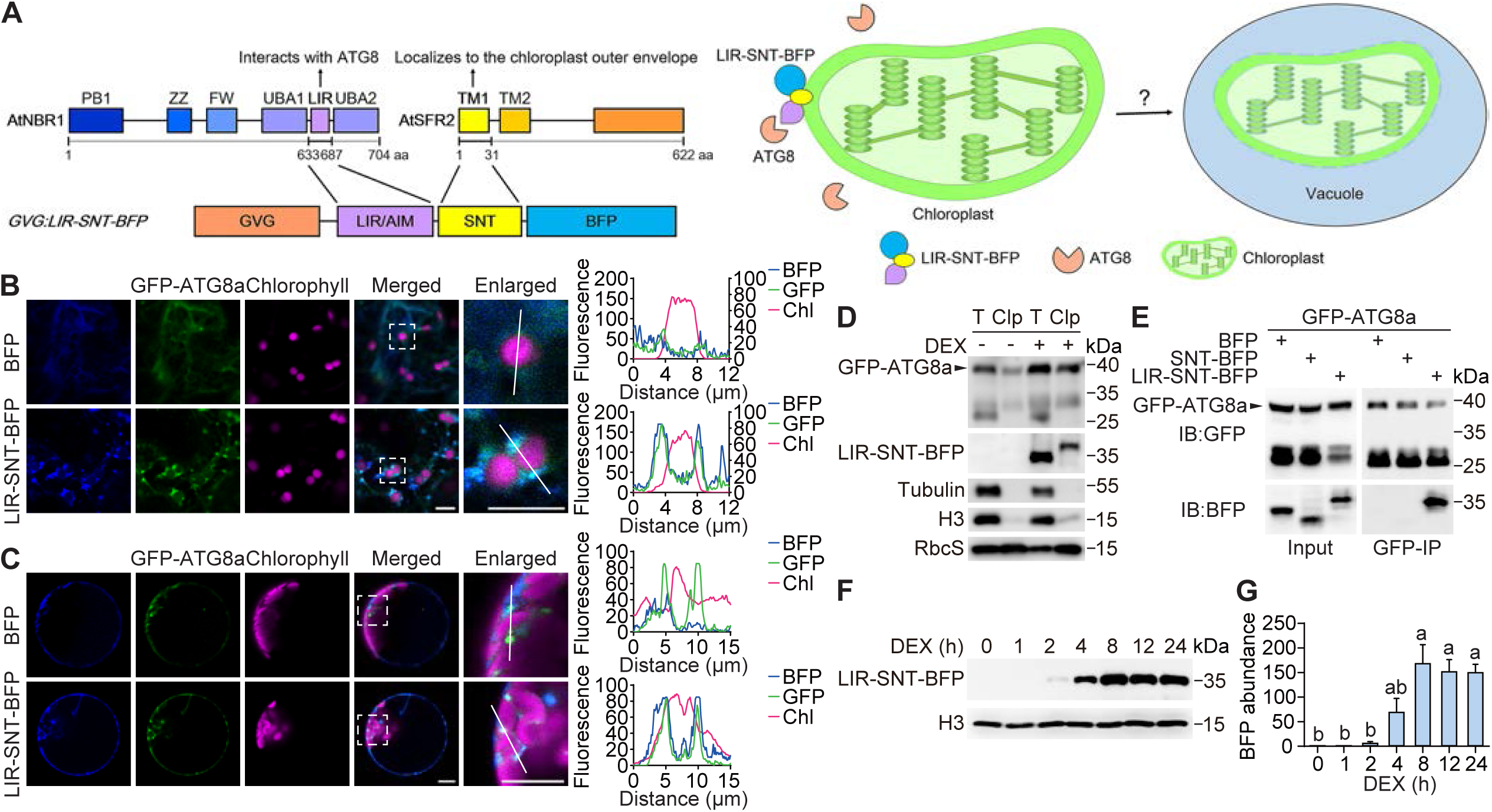
Development of a synthetic chloroplast autophagy receptor. **(A)** Diagram of the synthetic chlorophagy receptor. The synthetic receptor GVG:LIR-SNT-BFP comprises the LIR/AIM from AtNBR1 (LIR/AIM, aa633-687), the N-terminal amphipathic helix of AtSFR2 (SNT, aa1-31), and a fluorescent protein BFP, fused in frame after a glucocorticoid receptor-based inducible gene expression system (GVG). GVG:LIR-SNT-BFP is expected to localize to the outer envelope of chloroplasts, where it recruits ATG8 to facilitate autophagic degradation of chloroplasts, following Dexamethasone (DEX) treatment. **(B)** The synthetic chlorophagy receptor LIR-SNT-BFP coats chloroplasts and colocalizes with ATG8a in transiently transformed tobacco. *N.benthamiana* leaf epidermal cells transiently co-expressing *GFP-ATG8a* and *BFP* or *LIR-SNT-BFP* were analyzed by confocal microscopy. Bar = 10 μm. **(C)** LIR-SNT-BFP coats chloroplasts and colocalizes with ATG8a in transiently transformed Arabidopsis protoplast. Mesophyll protoplasts prepared from 4-week-old *GFP-ATG8a* plants were transiently transformed with BFP or *LIR-SNT-BFP* and analyzed by confocal microscopy. Bar = 10 μm. The colocalization was determined by calculating fluorescence intensity along the white lines in (B) and (C). **(D)** Western blot (WB) analysis of LIR-SNT-BFP and GFP-ATG8a on purified chloroplasts. Total proteins (T) and purified chloroplasts proteins (Clp) of the transgenic Arabidopsis expressing *GFP-ATG8a/LIR-SNT-BFP*, treated with DMSO or 30 μM DEX, were analyzed. Antibodies towards GFP, tRFP, Tubulin (cytoplasm marker), Histone H3 (H3, nucleus marker), and RubisCO small subunit (RbcS, chloroplast marker) were used. **(E)** Co-immunoprecipitation assay showing the *in vivo* interaction between GFP-ATG8a and LIR-SNT-BFP. Transgenic lines co-expressing *GFP-ATG8a* and *BFP*, *SNT-BFP*, or *LIR-SNT-BFP* were treated with 30 μM DEX before co-IP. Proteins were immunoprecipitated with GFP-Trap beads and detected with anti-GFP and anti-tRFP. **(F)** DEX-induced expression of LIR-SNT-BFP. Fourteen-day-old transgenic seedlings expressing *GFP-ATG8a/LIR-SNT-BFP* were treated with 30 μM DEX, harvested at indicated time points, and analyzed by WB. Anti-tRFP was used to detect LIR-SNT-BFP. Histone H3 (H3) served as an internal control. **(G)** Band intensities of LIR-SNT-BFP in (F) were quantified and normalized to H3. Data are means ± SEM of three biological replicates. Different letters denote statistically significant differences (*p* < 0.05) using Tukey’s honestly significant difference (HSD) test. Representative images or western blots from three biological replicates are shown in (B), (C), (D), (E), and (F).

We transformed an autophagy marker line, *ProUBQ10:GFP-ATG8a* (*GFP-ATG8a*) (Luo *et al*, 2017), with the synthetic chlorophagy receptor, and obtained T3 transgenic lines that carry both *GFP-ATG8a* and *GVG:LIR-SNT-BFP* (**Fig. S2A and B**). We also obtained T3 transgenic lines carrying *GFP-ATG8a* and *GVG:SNT-BFP*, which serve as a negative control (**Fig. S2A and B**). To preclude lines with T-DNA insertion in photosynthetic genes, T-DNA insertion sites in individual lines were determined by thermal asymmetric interlaced (TAIL)-PCR (**Fig. S2C and D**). Line 1, Line 2, and Line 27 were used in most experiments, for they have intergenic T-DNA insertions, and relatively stable DEX-induced expression of the receptor.

We found that BFP signals quench easily during confocal imaging in stable transgenic lines. Hence we took a biochemical approach to validate that the synthetic chlorophagy receptor can recruit ATG8 to the chloroplasts. Chloroplasts from the transgenic lines were purified with differential centrifugation and Percoll gradient centrifugation, and the presence of LIR-SNT-BFP and GFP-ATG8a on the purified chloroplasts after DEX induction were verified with western blotting (**Fig. 1D**). Notably, ATG8a was detected on the purified chloroplasts before DEX induction (**Fig. 1D**), which is in line with a previous report. (Wan *et al*, 2023) The interaction between ATG8a and the synthetic chlorophagy receptor was verified with co-immunoprecipitation (co-IP) (**Fig. 1E**). Neither BFP nor SNT-BFP, but LIR-SNT-BFP interacts with GFP-ATG8a in the double transgenic lines co-expressing the GFP-ATG8a and BFP-tagged proteins (**Fig. 1E**). The DEX-induced expression of the synthetic chlorophagy receptor starts from 4 h, and can be steadily observed after 8 h (**Fig. 1F and G**).

### The synthetic chlorophagy receptor facilitates autophagic degradation of chloroplast proteins

To see if and how the DEX-induced expression of the synthetic receptor may induce chloroplast autophagy, we examined the GFP signals in the mesophyll cells of mock and DEX-treated *GFP-ATG8a*/*LIR-SNT-BFP* seedlings. Seedlings carrying *GFP-ATG8a/SNT-BFP* served as a negative control. First, in line with the western blot results (**Fig. 1D**), and as reported before (Wan *et al*., 2023), GFP-ATG8a can be seen on the surface of chloroplasts (**Fig. 2A**). Treating *GFP-ATG8a/LIR-SNT-BFP* with DEX induced a 5-fold accumulation of GFP-ATG8a puncta on chloroplasts, which was not observed in *GFP-ATG8a/SNT-BFP* (**Fig. 2A and B**), validating the effective recruitment of ATG8 by the synthetic receptor. Consistently, by tracking the autophagic flux with GFP-ATG8 cleavage assay (**Fig. 2C**), we showed that autophagy was significantly induced upon LIR-SNT-BFP accumulation, but stayed unchanged in DEX-treated *GFP-ATG8a* or *GFP-ATG8a/SNT-BFP* seedlings (**Fig. 2C and D**).

**Fig 2.**
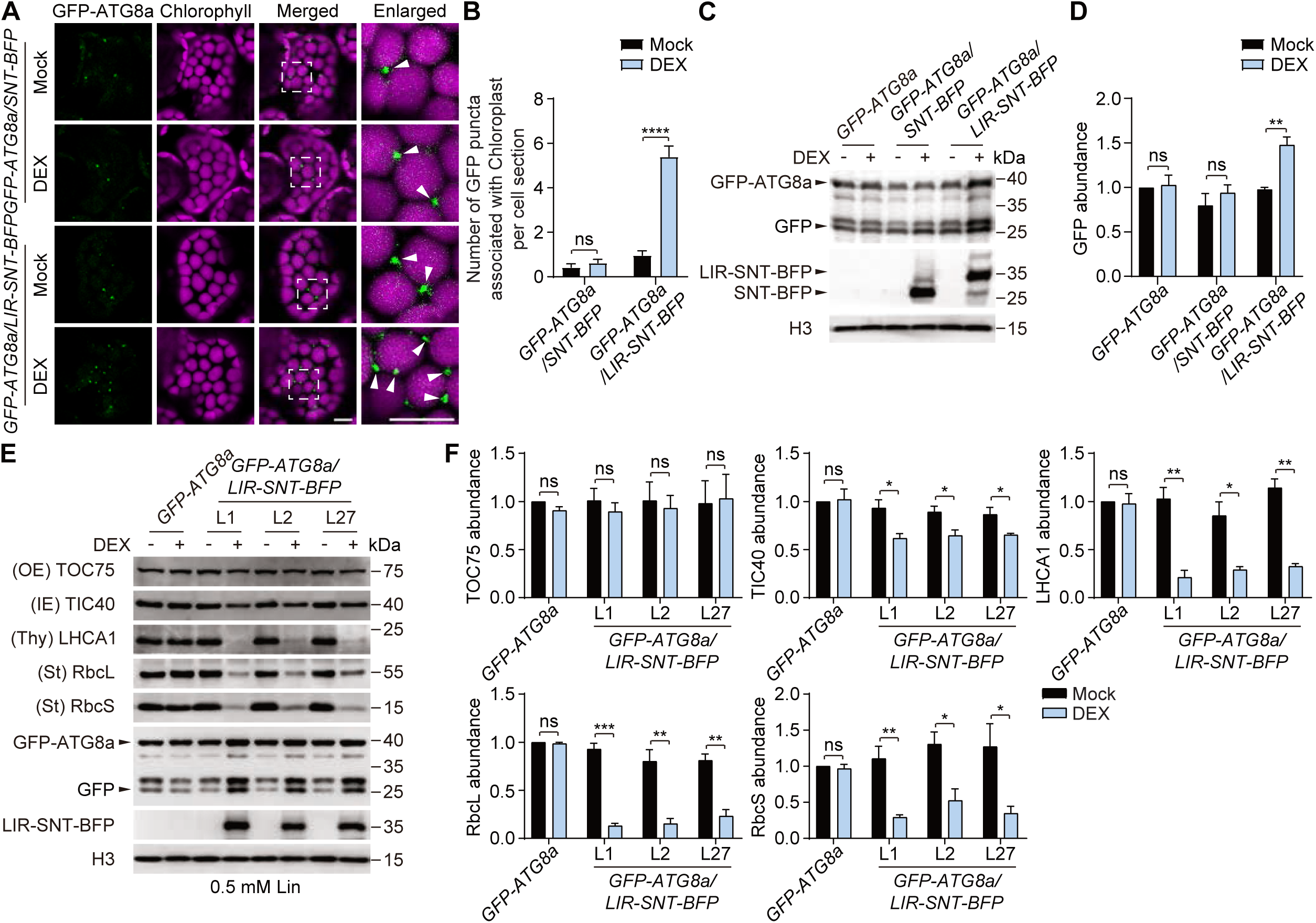
The synthetic chlorophagy receptor facilitates autophagic degradation of chloroplasts. **(A)** Expression of LIR-SNT-BFP but not SNT-BFP promotes GFP-ATG8a recruitment to the chloroplast. Five-day-old seedlings expressing *LIR-SNT-BFP* or *SNT-BFP* in the *GFP-ATG8a* background were treated with DMSO or 10 μM DEX for 3 days, before confocal imaging. White arrowheads indicate clusters of GFP-ATG8a on chloroplasts. Bar = 10 μm. **(B)** Quantification of the number of GFP puncta associated with chloroplasts per cell section in (A). Data are means ± SEM (n = 35). **(C)** Autophagy is induced by induction of LIR-SNT-BFP. Five-day-old seedlings expressing *LIR-SNT-BFP* or *SNT-BFP* in the *GFP-ATG8a* background were treated with DMSO or 10 μM DEX for 10 days before WB analysis of GFP-cleavage from GFP-ATG8a, indicative of autophagic flux. Anti-GFP and Anti-tRFP were used to detect GFP-ATG8a, free GFP, LIR-SNT-BFP and SNT-BFP. Histone H3 (H3) served as an internal control. **(D)** Band intensities of free GFP in (C) were quantified and normalized to H3. **(E)** LIR-SNT-BFP promotes the degradation of chloroplast proteins in the presence of lincomycin (Lin), an inhibitor of chloroplast protein synthesis. Five-day-old seedlings expressing *LIR-SNT-BFP* in the *GFP-ATG8a* background were treated with DMSO or 10 μM DEX plus 0.5 mM Lin for 14 days, before WB detection of chloroplast proteins TOC75 (chloroplast outer envelope protein), TIC40 (inner envelope protein), LHCA1 (thylakoid protein), RbcL (chloroplast-encoded stromal protein), and RbcS (stromal protein). Anti-GFP and Anti-tRFP were used to detect GFP-ATG8a and LIR-SNT-BFP. Histone H3 (H3) served as an internal control. **(F)** Band intensities in (E) were quantified and normalized to H3. *GFP-ATG8a* in DMSO (DEX-) was set as 100%. Data are means ± SEM of three biological replicates. ****, *p* < 0.0001; ***, *p* < 0.001; **, *p* < 0.01; *, *p* < 0.05; ns, no significant difference (student’s t-test). Representative images or western blots from three biological replicates are shown in (A), (C), (E). OE, chloroplast outer envelope; IE, chloroplast inner envelope; Thy, thylakoids; St, stroma.

To see if the chloroplasts were degraded as a consequence of induced autophagy, we compared the levels of chloroplast proteins that localize to the outer envelope (TOC75), inner envelope (TIC40), thylakoids (LHCA1), and stroma (RbcL and RbcS), before and after DEX induction. The levels of chloroplast proteins examined were almost unchanged after DEX induction (**Fig. S3A and B**). We reasoned that the degradation of chloroplast proteins could have been masked by new protein synthesis, hence treated the plants with Lincomycin, a lincosamide antibiotic that specifically inhibits chloroplast protein translation (Linnane & Stewart, 1967). Indeed, except for TOC75, whose level is reported to be regulated by NBR1-mediated selective autophagy (Wan *et al*., 2023), levels of chloroplast inner membrane proteins, thylakoid proteins, and stromal proteins were significantly reduced by synthetic receptor induction in the presence of Lincomycin (**Fig. 2E and F**).

To further validate that the amphipathic α-helix of SFR2 (SNT) can mediate chloroplast targeting and membrane binding of LIR, we generated an alternative synthetic receptor, *GVG:SNT-BFP-LIR*, by fusing LIR after, rather than before SNT, and obtained transgenic lines as for *GVG:LIR-SNT-BFP*. Same as the original receptor, this receptor interacted with ATG8a *in vivo* (**Fig. S4A**). The alternative receptor responded similarly to DEX induction and led to the degradation of LHCA1 and RbcL (**Fig. S4B-D**). These observations confirmed that LIR can be fused to either side of the amphipathic α-helix of SFR2 to elicit chlorophagy, although the C-terminal fusion of LIR appeared to be less effective.

### Moderate induction of the synthetic receptor promotes plant growth and partially protects leaves from herbicides

Previous studies have revealed the critical role of chlorophagy in protecting plants from adverse environmental conditions, such as prolonged carbon starvation, photodamage, heat, or UVB (Zhuang & Jiang, 2019). Nevertheless, it remains unknown if induced chlorophagy, in a certain range, can enhance plant growth. We took advantage of the inducible expression of the synthetic receptor and treated *GFP-ATG8a/LIR-SNT-BFP* lines with 0.01 to 10 µM DEX for 10 days to compare their rosette sizes. For Line 16, in which the synthetic receptor accumulated to a lower level (**Fig. S2A and B**), 0.1 to 10 µM DEX led to larger rosette. For Line 27, in which the synthetic receptor was highly expressed (**Fig. S2A and B**), 0.01 to 0.1 µM DEX treatments increased the rosette size, however 10 µM DEX treatment led to reduced rosette size (**Fig. 3A and B**). In addition, Line 27, treated with 0.1 to 10 µM DEX, had reduced chlorophyll contents (**Fig. 3C**). We examined the autophagic fluxes and the expression levels of the synthetic receptor in these plants. A positive correlation between the expression level of the synthetic receptor and the level of autophagy indicated by the increment of free GFP processed from GFP-ATG8a, was observed (**Fig. 3D-F**). We also found that, when the synthetic receptor is induced to a certain level range, which corresponds to 0.1 to 10 µM DEX in Line 16 and 0 to 0.01 µM DEX in Line 27, plant growth is promoted to a similar extent. When the synthetic receptor accumulated beyond this level, plant growth is inhibited, and the chlorophyll content drops (**Fig. 3A-F**). Clearly, in a certain range, plant growth correlates with the expression level of the synthetic chlorophagy receptor and the level of autophagy.

**Fig 3.**
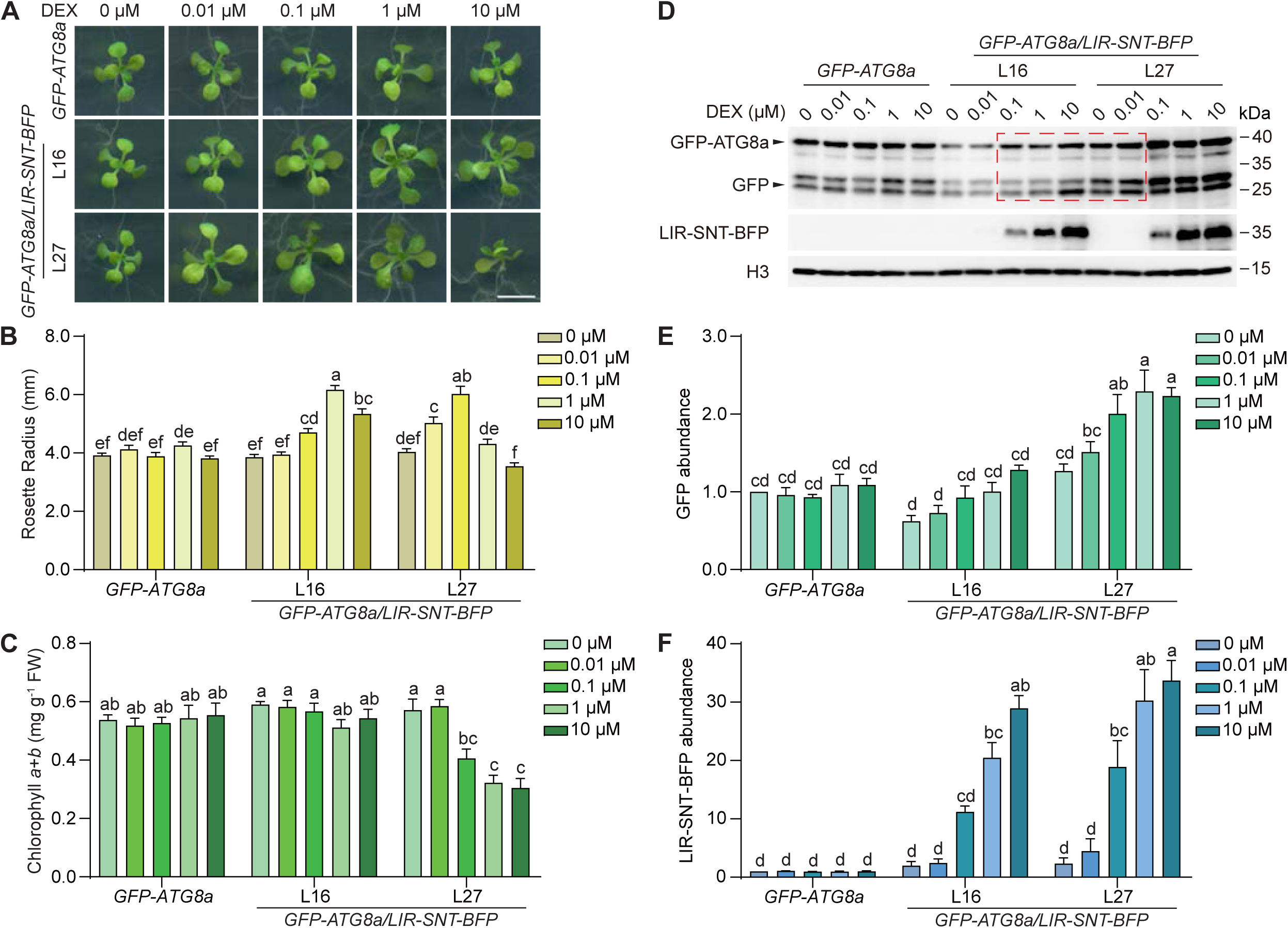
Moderate induction of the synthetic chlorophagy receptor promotes plant growth, whereas excessive induction inhibits plant growth. **(A)** Five-day-old seedlings expressing *LIR-SNT-BFP* in the *GFP-ATG8a* background were treated with indicated concentrations of DEX for 10 days. Bar = 5 mm. **(B)** Rosette radius measured from plants in (A). Data are means ± SEM (n = 16). **(C)** Chlorophyll contents of plants in (A). Data are means ± SEM (n = 20). **(D)** Autophagic flux increment roughly correlates with accumulation of LIR-SNT-BFP. Total proteins were extracted from plants in (A). Anti-GFP and Anti-tRFP were used to detect GFP-ATG8a, free GFP, and LIR-SNT-BFP. Histone H3 (H3) served as an internal control. **(E)** Band intensities of free GFP indicated the level of autophagic flux in (D) were quantified and normalized to H3. **(F)** Band intensities of LIR-SNT-BFP induced by DEX treatment in (D) were quantified and normalized to H3. *GFP-ATG8a* in DMSO (DEX-) was set as 100%. Data are means ± SEM of three biological replicates. Different letters indicate statistically significant differences (*p*< 0.05), as determined with a Tukey’s honestly significant difference (HSD) test. Representative images or western blots from three biological replicates are shown in (A) and (D).

We reasoned that artificially induced chlorophagy might accelerate the removal of damaged chloroplast components, thus might help alleviating the damage caused by certain herbicides. We tested *GFP-ATG8a/LIR-SNT-BFP* plants on Norflurazon (NF), a pyridazinone herbicide that disrupts carotenoid synthesis (Park *et al*, 2017), and 3-(3,4-dichlorophenyl)-1,1-dimethylurea (DCMU), an herbicide of the arylurea class that inhibits photosynthetic electron transport (Ridley, 1977). Both NF and DCMU led to photo-bleaching of cotyledons in *GFP-ATG8a* and *GFP-ATG8a/LIR-SNT-BFP*. In contrast, DEX induction of the synthetic chlorophagy receptor partially reduced the chlorosis (**Fig. 4A and B**). Confocal microscopy confirmed the reduced cell death, indicated by propidium iodide (PI) staining, and reduced chlorophyll loss caused by NF or DCMU in DEX-treated plants expressing *GFP-ATG8a/LIR-SNT-BFP* (**Fig. 4C**), but not in the plants expressing *GFP-ATG8a* (**Fig. S5A**). Particularly, the uneven distribution of chlorophyll caused by NF or DCMU, indicative of defective chloroplast structure and function, was partially restored in DEX-treated plants that expressed the synthetic chlorophagy receptor (**Fig. 4D**), but not in the control line *GFP-ATG8a* (**Fig. S5B**). In seedlings grown in soil, we also observed partial suppression of NF-induced plant death by induced expression of the synthetic chlorophagy receptor (**Fig. S5C**). We concluded that, artificially-induced chloroplast autophagy, which likely exceeds the level of endogenous chlorophagy, can help maintain chloroplast homeostasis when chlorophyll synthesis or electron transport is compromised by certain herbicides.

**Fig 4.**
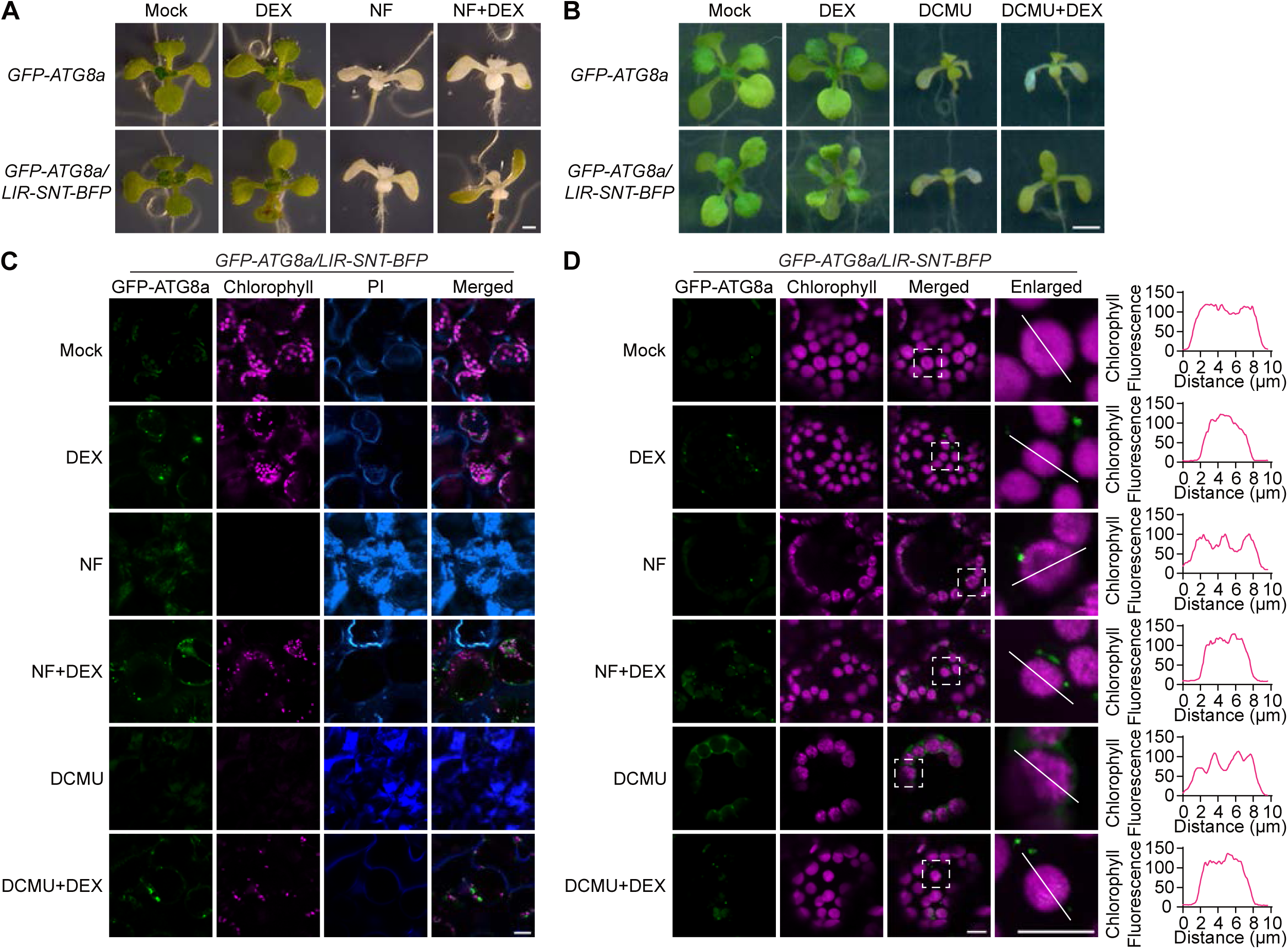
Induction of the synthetic chlorophagy receptor partially protects plant from herbicides norflurazon and DCMU. **(A)** Norflurazon (NF) led to leaf chlorosis, whereas LIR-SNT-BFP induction partially protected cotyledons from chlorosis. Five-day-old seedlings expressing *LIR-SNT-BFP* in the *GFP-ATG8a* background were treated with 10 μM DEX, or 1 μM NF, or DEX plus NF for 7 days. Bar = 1 mm. **(B)** DCMU led to leaf chlorosis, whereas LIR-SNT-BFP induction partially protected cotyledons from chlorosis. Five-day-old seedlings expressing *LIR-SNT-BFP* in the *GFP-ATG8a* background were treated with 10 μM DEX, or 2 μM DCMU, or DEX plus DCMU for 10 days. Bar = 2 mm. **(C)** NF- or DCMU-induced chlorophyll loss and cell death was partially suppressed by LIR-SNT-BFP induction. Five-day-old seedlings expressing *GFP-ATG8a/LIR-SNT-BFP* were treated with 10 μM DEX, 1 μM NF, 2 μM DCMU, or combinations of the chemicals as indicated for 10 days, before confocal imaging. Cell death was visualized by staining with propidium iodine (PI). Bar = 20 μm. **(D)** NF- or DCMU-induced abnormal, uneven distribution of chlorophyll was partially recovered by LIR-SNT-BFP induction. Five-day-old seedlings expressing *GFP-ATG8a/LIR-SNT-BFP* were treated with 10 μM DEX, 1 μM NF, 10 μM DCMU, or combinations of the chemicals as indicated for 3 days, before confocal imaging. Chlorophyll fluorescence along the white lines plotted to the right. Bar = 10 μm. Representative images from three biological replicates are shown in (A), (B), (C), and (D).

### Induction of the synthetic receptor promotes chloroplast division

In animals and yeasts, long tubular mitochondria divide before they can be engulfed by autophagosome (Frank *et al*, 2012; Fukuda *et al*, 2023). Chloroplasts are much larger than mitochondria, and various forms of chloroplast piecemeal autophagy, as well as microautophagy of the entire chloroplasts, have been reported (Izumi *et al*., 2024; Zhuang & Jiang, 2019). Yet still, the relationship between chloroplast division and autophagy is largely unknown. Since the chloroplasts became smaller in DEX-treated *GFP-ATG8a/LIR-SNT-BFP* plants (**Fig. 4D**), we asked if chloroplasts divide before they are degraded in the vacuole. Following the methods in chloroplast division studies (Okazaki *et al*, 2009b; Osteryoung *et al*, 1998), we quantified the numbers of chloroplasts per mesophyll cell in mock- and DEX-treated *GFP-ATG8a/LIR-SNT-BFP* plants, and saw significant increments in all DEX-treated lines that could be a consequence of chloroplast division (**Fig. 5A and B**). Indeed, we observed more dumbbell-shaped chloroplasts that appear to be undergoing fission in DEX-treated *GFP-ATG8a/LIR-SNT-BFP* mesophyll cells (**Fig. 5C and D**). Then, to see if known machineries mediate the division, we examined the protein expression level of PLASTID DIVISION2 (PDV2), in transgenic plants. PDV2 is a land plant specific integral outer envelope membrane protein that has a positive dosage effect on chloroplast division (Chang *et al*, 2017; Okazaki *et al*, 2009a). Mechanistically, PDV2 recruits the cytosolic dynamin-related protein DRP5B/ARC5 to the division site to complete chloroplast division (Miyagishima *et al*, 2006; Okazaki *et al*., 2009a). To our surprise, despite the increased chloroplast numbers, the protein level of PDV2 stayed unchanged in DEX-treated *GFP-ATG8a/LIR-SNT-BFP* plants (**Fig. 5E and F**). We then crossed the *pdv2* mutant into *GFP-ATG8a/LIR-SNT-BFP*. Confocal imaging and TEM showed that the giant chloroplast phenotype in *pdv2/GFP-ATG8a/LIR-SNT-BFP* stayed unchanged after DEX treatment (**Fig. 5G-H**). On the other hand, DEX-induced chloroplast protein degradation in *pdv2/GFP-ATG8a/LIR-SNT-BFP* was similar to what we observed in *GFP-ATG8a/LIR-SNT-BFP* (**Fig. 5I and J**). Likewise, the synthetic chlorophagy receptor partially protected *pdv2* from norflurazon (**Fig. S6A and B**). These results indicated that the giant chloroplasts in *pdv2* could still undergo synthetic receptor-elicited chlorophagy. Therefore, PDV2-mediated chloroplast division seems not required for this specific form of chlorophagy.

**Fig 5.**
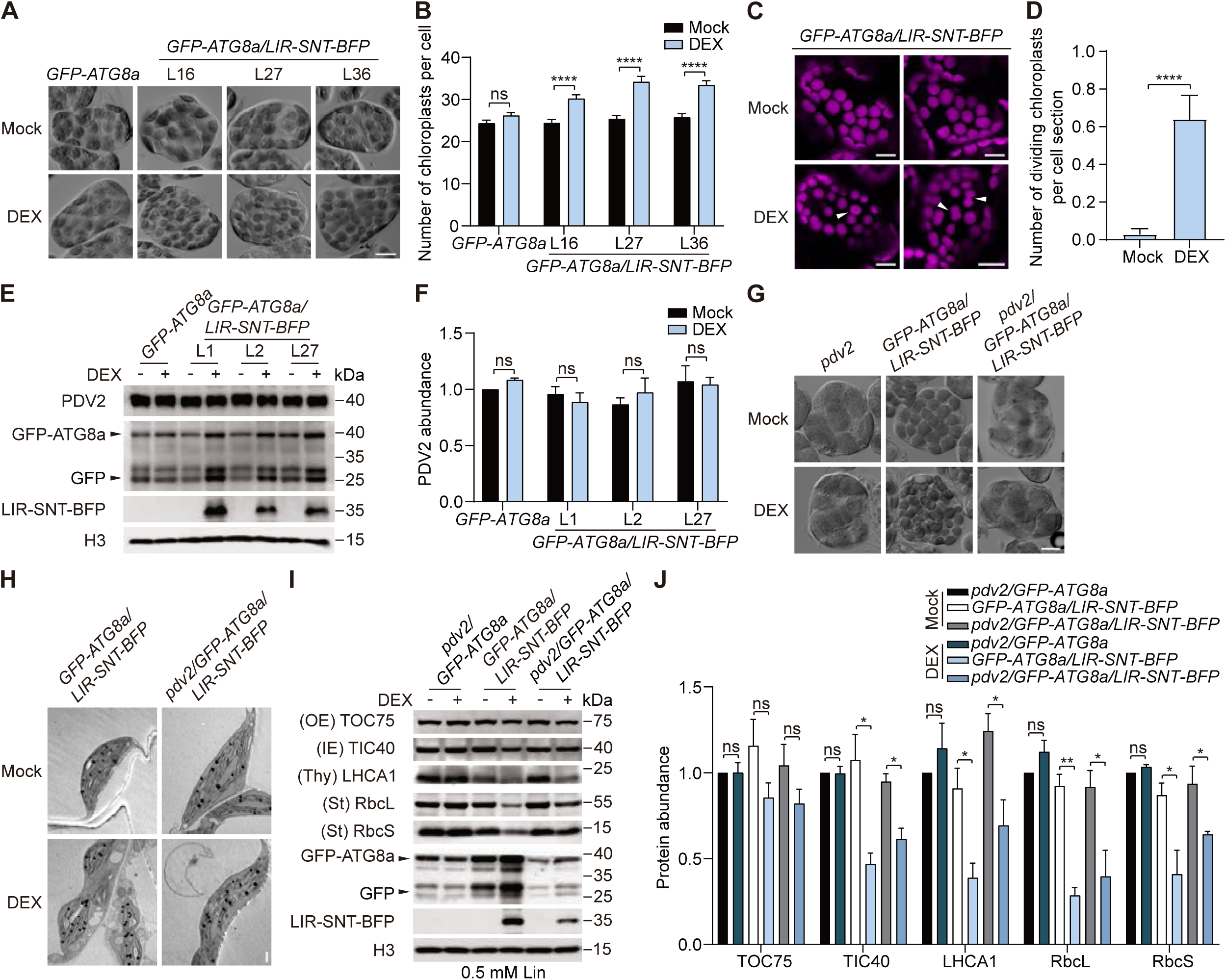
Induction of the synthetic chlorophagy receptor stimulates chloroplast division. **(A)** Induction of LIR-SNT-BFP significantly increases the number of chloroplasts per cell. Five-day-old seedlings expressing *LIR-SNT-BFP* in the *GFP-ATG8a* background were treated with DMSO or 10 μM DEX for 10 days. Mesophyll cells from the first pair of true leaves were fixed before confocal imaging. Bar = 10 μm. **(B)** Quantification of the number of chloroplasts per cell in (A). Data are means ± SEM (n = 30). **(C)** Induction of LIR-SNT-BFP significantly increases the incidence of chloroplast division per cell section. Five-day-old seedlings expressing *GFP-ATG8a/LIR-SNT-BFP* were treated with DMSO or 10 μM DEX for 5 days before confocal imaging. White arrowheads indicate dividing chloroplasts. Bar = 10 μm. **(D)** Quantification of the number of dividing chloroplasts per cell section in (C). Data are means ± SEM (n = 36). **(E)** PDV2 protein levels stayed unchanged before and after LIR-SNT-BFP induction. Five-day-old seedlings expressing *LIR-SNT-BFP* in the *GFP-ATG8a* background were treated with DMSO or 10 μM DEX for 10 days before WB analysis. Proteins were detected by anti-PDV2, anti-GFP, and anti-tRFP. Histone H3 (H3) served as an internal control. **(F)** Band intensities of PDV2 in (E) were quantified and normalized to H3. *GFP-ATG8a* in DMSO (DEX-) was set as 100%. **(G)** LIR-SNT-BFP induction did not change the giant chloroplast phenotype in *pdv2*. Five-day-old *pdv2*, *GFP-ATG8a/LIR-SNT-BFP*, and *pdv2/GFP-ATG8a/LIR-SNT-BFP* were treated with DMSO or 10 μM DEX for 10 days. Mesophyll cells from the first pair of true leaves were fixed before confocal imaging. Bar = 10 μm. **(H)** LIR-SNT-BFP induction did not change the giant chloroplast phenotype in *pdv2*. Cotyledons of five-day-old seedlings expressing *GFP-ATG8a/LIR-SNT-BFP* in the *pdv2* background, treated with DMSO or 10 μM DEX for 6 days, were analyzed by TEM. Bar = 1 μm. **(I)** LIR-SNT-BFP induction led to the degradation of chloroplast proteins in *pdv2*. Five-day-old *pdv2/GFP-ATG8a*, *LIR-SNT-BFP*/*GFP-ATG8a*, and *pdv2/GFP-ATG8a/LIR-SNT-BFP* were treated with DMSO or 10 μM DEX in the presence of 0.5 mM Lin for 14 days before WB detection of chloroplast proteins TOC75 (chloroplast outer envelope protein), TIC40 (inner envelope protein), LHCA1 (thylakoid protein), RbcL (chloroplast encoded stromal protein), and RbcS (stromal protein). Anti-GFP and Anti-tRFP were used to detect GFP-ATG8a and LIR-SNT-BFP. Histone H3 (H3) served as an internal control. **(J)** Band intensities in (I) were quantified and normalized to H3. *pdv2/GFP-ATG8a* in DMSO (DEX-) was set as 100%. Data are means ± SEM of three biological replicates. ****, *p* < 0.0001; ***, *p* < 0.001; **, *p* < 0.01; *, *p* < 0.05; ns, no significant difference (student’s t-test). Representative images or western blots from three biological replicates are shown in (A), (C), (E), (G), (H), and (I). OE, chloroplast outer envelope; IE, chloroplast inner envelope; Thy, thylakoids; St, stroma.

### The synthetic receptor-elicited chlorophagy does not require the ATG8 conjugation machinery proteins ATG5 and ATG7

To gain more insights into the chloroplast autophagy elicited by the synthetic receptor, we crossed three autophagy-deficient mutants, *atg2*, *atg5*, and *atg7*, into *GFP-ATG8a/LIR-SNT-BFP*. ATG2 is a lipid transfer protein key to phagophore expansion and autophagosome closure (Luo *et al*, 2023a), ATG5 and ATG7 are required for ATG8 conjugation (Thompson *et al*, 2005). We obtained *atg5/GFP-ATG8a/LIR-SNT-BFP* and *atg7/GFP-ATG8a/LIR-SNT-BFP* lines, but failed to recover lines that can express the synthetic receptor in the *atg2* background (**Fig. S7A-C**). Interestingly, DEX-induced synthetic receptor partially rescued the photo-bleaching and uneven distribution of chlorophyll in norflurazon-treated *atg5/GFP-ATG8a/LIR-SNT-BFP* and *atg7/GFP-ATG8a/LIR-SNT-BFP*, like in *GFP-ATG8a/LIR-SNT-BFP* (**Fig. 6A and B**). Consistent with this, DEX-induced chloroplast protein degradation in *atg5* and *atg7* in the presence of lincomycin was comparable to the wild-type background (**Fig. 6C and D**). Reduced chloroplast sizes upon DEX treatment were also observed in *atg5* and *atg7* like in the wild-type background (**Fig. S7D and E**). All these observations indicated that ATG5 and ATG7, key components of the ATG8 conjugation machinery, are dispensable for this specific form of chloroplast autophagy.

**Fig 6.**
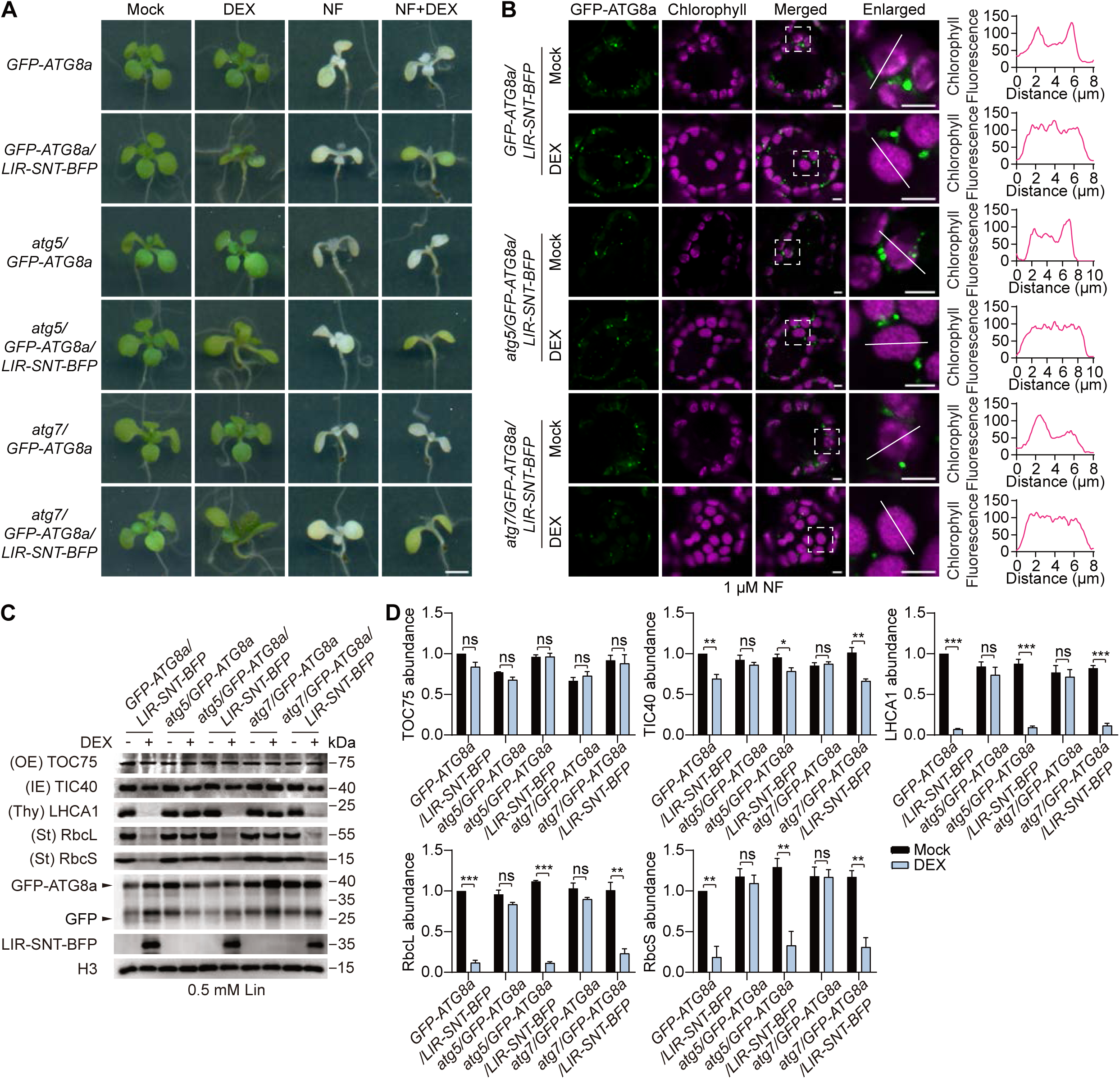
The synthetic receptor-elicited chlorophagy does not require the ATG8 conjugation machinery proteins ATG5 and ATG7. **(A)** LIR-SNT-BFP induction partially protected cotyledons from chlorosis in NF-treated *atg5* and *atg7*. Five-day-old seedlings expressing *GFP-ATG8a* or *GFP-ATG8a/LIR-SNT-BFP* in the *atg5* or *atg7* background were treated with 10 μM DEX, or 1 μM NF, or NF plus DEX for 7 days. Bar = 2 mm. **(B)** LIR-SNT-BFP induction led to partial recovery of the abnormal, uneven distribution of chlorophyll in NF-treated *atg5* and *atg7*. Five-day-old seedlings expressing *GFP-ATG8a/LIR-SNT-BFP* in the *atg5* or *atg7* background were treated with 1 μM NF, with or without 10 μM DEX for 3 days, before confocal imaging. Chlorophyll fluorescence along the white lines plotted to the right. Bar = 5 μm. **(C)** LIR-SNT-BFP induction promotes the degradation of chloroplast proteins in *atg5* and *atg7*. Five-day-old seedlings expressing *GFP-ATG8a* or *GFP-ATG8a/LIR-SNT-BFP* in the *atg5* or *atg7* background were treated with DMSO or 10 μM DEX in the presence of 0.5 mM Lin for 14 days before WB detection of chloroplast proteins TOC75 (chloroplast outer envelope protein), TIC40 (inner envelope protein), LHCA1 (thylakoid protein), RbcL (chloroplast encoded stromal protein), and RbcS (stromal protein). Anti-GFP and Anti-tRFP were used to detect GFP-ATG8a and LIR-SNT-BFP. Histone H3 (H3) served as an internal control. **(D)** Band intensities in (C) were quantified and normalized to H3. *GFP-ATG8a/LIR-SNT-BFP* in DMSO (DEX-) was set as 100%. Data are means ± SEM of three biological replicates. ***, *p* < 0.001; **, *p* < 0.01; *, *p* < 0.05; ns, no significant difference (student’s t-test). Representative images or western blots from three biological replicates are shown in (A), (B), and (C). OE, chloroplast outer envelope; IE, chloroplast inner envelope; Thy, thylakoids; St, stroma.

### The synthetic receptor-elicited chlorophagy is likely a form of microautophagy

We speculated that the ATG5- and ATG7-independent synthetic receptor-mediated chlorophagy may be a form of microautophagy with the lytic vacuoles devouring the chloroplasts. To explore this possibility, we introduced a tonoplast marker, *YFP-VAMP711* (Geldner *et al*, 2009), into *GFP-ATG8a/LIR-SNT-BFP* by crossing, and examined the relationship between chloroplasts and vacuoles with confocal imaging. Indeed, tonoplast invaginations surrounding individual chloroplasts increased by approximately 2-fold after DEX induction in mesophyll cells and in protoplasts prepared from this triple transgenic line (**Fig. 7A-D**). We then employed live-cell imaging to capture the dynamic interaction between the chloroplasts and the vacuoles. Without DEX-induction of the synthetic chlorophagy receptor, the tonoplast surrounding the chloroplasts was relatively quiet, with sporadic trans-strands observed (**Fig. 7E, Movies 1 and 2**). After DEX-induced expression of LIR-SNT-BFP, however, the tonoplast became highly dynamic and actively chased chloroplasts and wrapped around them (**Fig. 7E, Movies 3 and 4**). Chloroplasts wrapped in vacuoles were frequently observed after DEX treatment, and small vacuoles were often seen beside the chloroplasts, likely resulting from vacuole fission after invagination of chloroplasts (**Fig. 7F, Movie 4**).

**Fig 7.**
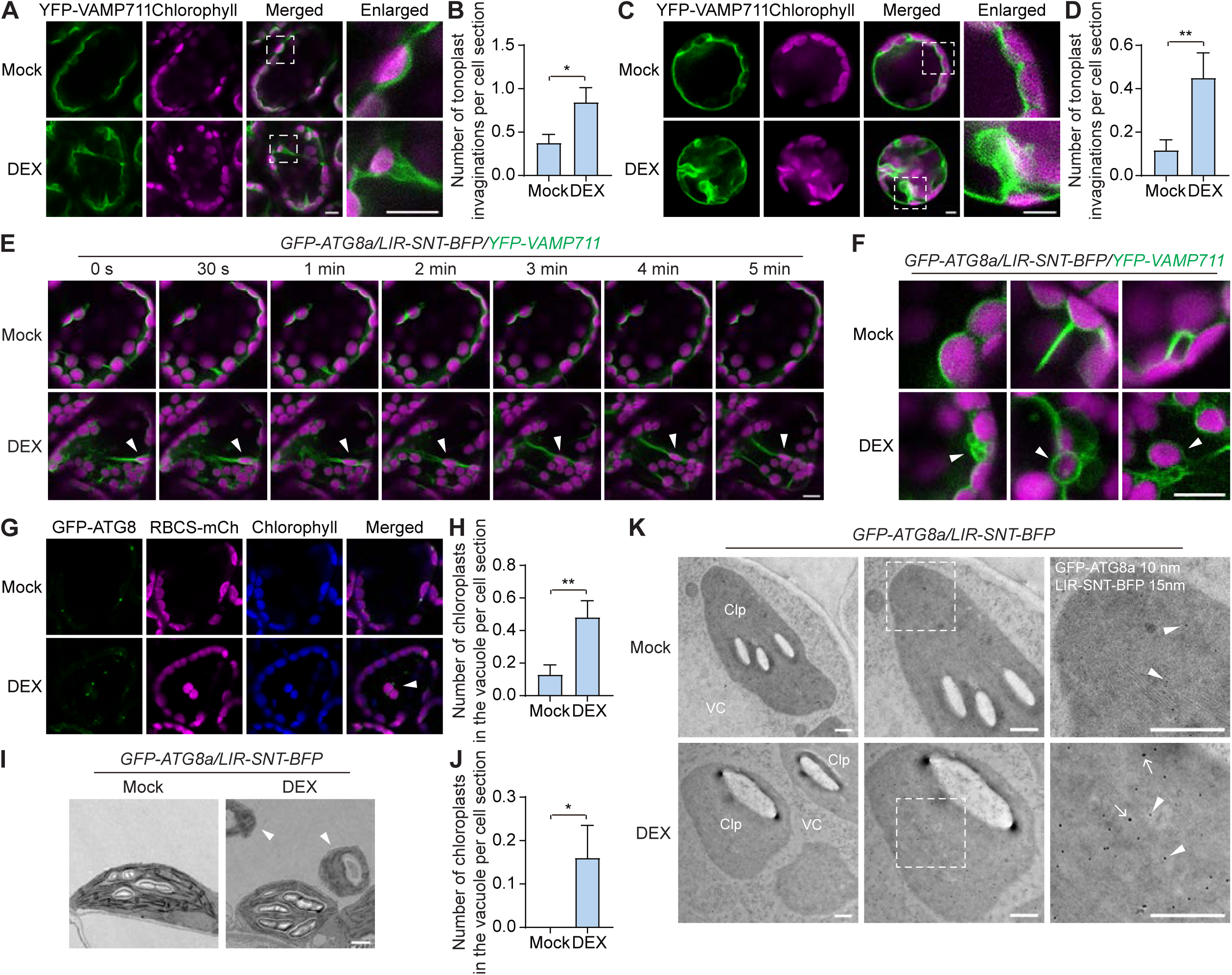
The synthetic receptor-elicited chlorophagy is likely a form of microautophagy. **(A)** Tonoplast invaginations that wrap chloroplasts were observed after LIR-SNT-BFP induction. Five-day-old seedlings expressing *GFP-ATG8a/LIR-SNT-BFP/YFP-VAMP711* were treated with DMSO or 10 μM DEX for 5 days before confocal imaging. Bar = 10 μm. **(B)** Quantification of the number of tonoplast invaginations per cell section in (A). Data are means ± SEM (n = 32). **(C)** Tonoplast invaginations that wrap chloroplasts were frequently observed after LIR-SNT-BFP induction in protoplasts. Fourteen-day-old seedlings expressing *GFP-ATG8a/LIR-SNT-BFP/YFP-VAMP711*, were treated with DMSO or 30 μM DEX for 12 hours, before preparation of protoplasts. Bar = 5 μm. **(D)** Quantification of the number of tonoplast invaginations per cell section in (C). Data are means ± SEM (n = 60). **(E)** Real-time imaging of tonoplast invagination that wraps chloroplasts (white arrowheads) after LIR-SNT-BFP induction. Five-day-old seedlings expressing *GFP-ATG8a/LIR-SNT-BFP/YFP-VAMP711* were treated with DMSO or 10 μM DEX for 5 days before live-cell imaging. Bar = 10 μm. **(F)** Chloroplasts wrapped by vacuoles and clusters of small vacuoles (white arrowheads) were observed after LIR-SNT-BFP induction. Cotyledons of *GFP-ATG8a/LIR-SNT-BFP/YFP-VAMP711* plants in (E) were analyzed by confocal imaging. Bar = 10 μm. **(G)** Chloroplasts are present in the central vacuole after LIR-SNT-BFP induction. Five-day-old seedlings expressing *GFP-ATG8a/LIR-SNT-BFP/RBCS1A-mCherry* were treated with DMSO or 10 μM DEX for 5 days. The V-ATPase inhibitor Concanamycin A (ConA) (1 μM) was added 24 hours before confocal imaging. White arrowhead indicates chloroplasts in the vacuole. Bar = 10 μm. **(H)** Quantification of the number of chloroplasts in the vacuole per cell section in (G). Data are means ± SEM (n = 54). **(I)** Presence of chloroplasts in the central vacuole after LIR-SNT-BFP induction revealed by TEM. Five-day-old seedlings expressing *GFP-ATG8a/LIR-SNT-BFP* were treated with DMSO or 10 μM DEX for 5 days, and the cotyledons were chemically fixed and analyzed by transmission electron microscopy (TEM). White arrowheads indicate the chloroplasts in the vacuole. Bar = 1 μm. **(J)** Quantification of the number of chloroplasts in the vacuole per image in (I). Data are means ± SEM (n = 25). **(K)** Detection of LIR-SNT-BFP and GFP-ATG8a on the chloroplasts in the vacuole by immuno-EM (IEM). Five-day-old seedlings expressing *GFP-ATG8a/LIR-SNT-BFP* were treated with DMSO or 10 μM DEX for 3 days, with 1 μM ConA added 24 hours before fixation. The cotyledons were chemically fixed and analyzed by IEM. The samples were double immunolabeled with anti-GFP (arrowheads) followed by 10-nm gold-conjugated goat anti-mouse IgG, and anti-BFP (arrows) followed by 15-nm gold-conjugated goat anti-rabbit IgG. Bar = 0.5 μm. **, *p* < 0.01; *, *p* < 0.05 (student’s t-test). Representative images from three biological replicates are shown in (A), (C), (E), (F), (G), (I), and (K). Clp, chloroplast; VC, vacuole.

We also generated stable transgenic lines carrying *RBCS1A-mCherry/GFP-ATG8a/LIR-SNT-BFP* to document chloroplast degradation by looking at RBCS1A-mCherry. We found that entire chloroplasts, instead of mCherry-labelled puncta that represent RCB, accumulated in the vacuole after DEX-induction of LIR-SNT-BFP (**Fig 7G and H**). Similarly, TEM analyses showed that the number of chloroplasts in the vacuoles largely increased in the mesophyll cells of *GFP-ATG8a/LIR-SNT-BFP* after DEX treatment (**Fig. 7I and J**). We further performed double-label immuno-electron microscopy on the mesophyll cells of mock- or DEX-treated *GFP-ATG8a/LIR-SNT-BFP* seedlings, and examined whether entire chloroplasts in the vacuole observed after DEX treatment are synthetic receptor-positive. GFP-ATG8a signals, labelled by secondary antibody conjugated to gold particles 10 nm in diameter, were sporadically seen on the chloroplasts in mock-treated *GFP-ATG8a/LIR-SNT-BFP*. In contrast, in the mesophyll cells of DEX-treated *GFP-ATG8a/LIR-SNT-BFP*, chloroplasts in the vacuole were labelled with both the synthetic receptor LIR-SNT-BFP (larger gold particles, 15 nm in diameter) and GFP-ATG8a (smaller gold particles) (**Fig. 7K**). These results suggested that the chloroplasts were indeed degraded in the vacuole by microautophagy elicited by the synthetic receptor.

## DISCUSSION

### Inducible degradation of whole chloroplasts with an artificial/synthetic receptor: proof of concept and application potential

Even with over two decades of studies on plant autophagy, a canonical chloroplast autophagy receptor that mediates vacuolar degradation of the entire chloroplasts remains unidentified. As a proof of concept, we designed and constructed a chloroplast autophagy receptor that combines outer envelope localization, the ability to recruit ATG8, and a fluorescent protein tag (**Fig. 1, Fig. S1**). With an inducible expression system, this synthetic chlorophagy receptor accumulates only upon DEX treatment, thus may avoid silencing or lethal effects often associated with constitutively elevated chlorophagy. Induced expression of this synthetic receptor simultaneously promotes chloroplast autophagy and division (**Fig. 2**, **Fig. 5**). Surprisingly, it was a form of microautophagy that we observed, as vacuoles clearly engulfed the chloroplasts (**Fig. 7, Movies 1-4**). Neither ATG5 nor ATG7 is required for such microautophagy (**Fig. 6, Fig. S7**). Also to our surprise, although chloroplast division evidently accompanies chlorophagy, PDV2, a central organizer of chloroplast division machinery on the cytoplasmic side, is not required for this form of chlorophagy (**Fig. 5**). This is similar to carbon starvation-induced chloroplast piecemeal autophagy, in which DRP5B/ARC5, a dynamin-related protein recruited by PDV2 that constitutes the division ring on the cytoplasmic side(Liu *et al*, 2024), is not required(Izumi *et al*., 2024). However, the synthetic receptor-mediated chlorophagy is different from the piecemeal degradation of chloroplasts, for no RCB puncta were detected in the vacuole after the induction of the receptor (**Fig. 7**), and that the piecemeal chlorophagy does require ATG5 and ATG7(Izumi *et al*., 2024). Hence, both shared and different chloroplast autophagy mechanisms are involved between the two forms of chlorophagy.

Enhanced organelle autophagy, especially mitophagy, is critical to organismal homeostasis, increased fitness, and longevity in animal models(Chen *et al*, 2020; Klionsky *et al*, 2021). Nevertheless, the benefit of organelle autophagy induction has not been fully explored in plants. Using the synthetic chlorophagy receptor, we found out that increased chloroplast autophagy can promote plant growth, and that a certain upper level limit for autophagy in promoting plant growth exists, regardless of which transgenic line is examined (**Fig. 3**). Beyond this level, autophagy becomes detrimental and inhibits rosette growth. This to our knowledge is the first report on an upper level limit of plant autophagic flux with respect to optimal growth. Another interesting finding was the partial rescue of herbicide-induced chlorophyll damage and leaf chlorosis with induced chlorophagy (**Fig. 4**). Induced chlorophagy helped alleviating the damage on chlorophyll biosynthesis or electron transport, likely contributing to the homeostasis of the chloroplast population in a mesophyll cell. However, the tolerance to herbicides conferred by artificially induced chlorophagy also appeared to be limited.

### What really happens during chloroplast microautophagy?

So far, no canonical chlorophagy receptor has been reported. This synthetic chlorophagy we developed may partially mimic the yet unidentified endogenous chlorophagy receptor. We noticed several interesting facts during this specific form of chlorophagy. First, a small population of ATG8 already decorates chloroplast under control condition, as reported(Wan *et al*., 2023). Second, the chlorophagy we observed does not require the ATG8 conjugation machinery. Third, chloroplasts engulfed by vacuoles were observed when the synthetic receptor is induced. Also, we did not see protrusions that shed off chloroplasts like in carbon-starved leaves(Izumi *et al*., 2024). We speculated that both endogenous piecemeal autophagy, mediated by PE-conjugated ATG8, and microautophagy of the entire chloroplasts, may have taken place. The dramatic change in vacuole shape around the chloroplast is somewhat similar to early observation on peroxisome degradation by microautophagy(Sakai *et al*, 1998). When shifted from methanol medium to glucose medium, *Pichia pastoris* (*Komagataella phaffii*) and *Hansenula polymorpha*, two methylotrophic yeast species, would promptly degrade their enlarged peroxisomes that had been metabolizing methanol as a carbon source. The large size of chloroplasts and vacuoles can be compared to the large size of peroxisomes and vacuoles in the two yeast species, and some mechanisms may be shared between these forms of microautophagy. For instance, the ATG8 cluster on chloroplast may represent a structure similar to micropexophagy-specific membrane apparatus (MIPA) in *P.pastoris*, and the clustered vacuoles may be similar to vacuolar sequestering membrane (VSM) that facilitate microautophagy of large organelles. The detailed molecular mechanism, however, remains to be explored.

In summary, targeted degradation of a large organelle—the chloroplast, is achieved with an inducible synthetic autophagy receptor. The beneficial effects of moderately induced chloroplast autophagy included increased rosette size and partial protection against herbicides NF and DCMU. Both are potentially useful for crop breeding. With this tool we also uncovered interesting behaviors of chloroplasts and vacuoles, such as PDV2-independent chlorophagy, and microautophagy that appears independent of ATG8 lipidation.

## MATERIALS AND METHODS

### Plant materials and growth conditions

*Arabidopsis thaliana* used in this study was in Columbia-0 (Col-0) background. The T-DNA insertion mutants of *atg2-1* (SALK_076727), *atg5-1* (SAIL_129B07, CS806267), and *atg7* (SAIL_11H07, CS862226), were obtained from ABRC. *pdv2-1* (SALK_059656) was a gift from Cheng Chen (Shanghai Jiao Tong University). All the other plant materials used in this study were generated by crossing or floral dipping. Surface-sterilized Arabidopsis seeds were stratified at 4°C for 2 days in the dark. Transgenic plants expressing *GFP-ATG8a* were selected on half-strength Murashige and Skoog (1/2 MS) medium containing 20 mg/mL Hygromycin B. Transgenic plants co-expressing *LIR-SNT-BFP* and *GFP-ATG8a* were selected on 1/2 MS medium containing 20 mg/mL Hygromycin B and Basta (1:10000 dilution). Transgenic plants expressing *GFP-ATG8a/LIR-SNT-BFP/RBCS1A-mCherry* were selected on 1/2 MS medium containing 20 mg/mL Hygromycin B, 50 mg/mL Kana and Basta (1:10000 dilution).For DEX treatment, five-day-old seedlings were transferred to 1/2 MS medium supplemented with DMSO or 10 μM DEX. Plants were grown at 16 h (22°C)/8 h (18°C) with full spectrum LED lamps at 100 μE m^−2^ s^−1^. Primers used for genotyping are listed in **Table S1**.

### Plasmids construction

The synthetic chlorophagy receptor LIR-SNT-BFP contains the LIR domain of AtNBR1 (LIR, aa633-687) to interact with ATG8, the N-terminal of AtSFR2 (SNT, aa1-31) to localize to the chloroplast outer envelope, and a blue fluorescent protein BFP. For inducible expression of the receptor, a *pCAMBIA3301* vector carrying *GVG* (a glucocorticoid receptor-based inducible gene expression system) was constructed. The *GVG* gene is composed of the *CaMV 35S* promoter, *GAL4*-binding domain-*VP16* activation domain-*GR* fusion, the *rbcS-E9* terminator and upstream activation sequence (*UAS*), which were amplified from the *pTA7002* vector. Then the *LIR-SNT-BFP* fusion gene was inserted between Nco I and Eco91 I of *pCAMBIA3301*, in frame after *GVG* gene. Primers used for plasmid construction are listed in **Table S1**.

### Transient expression assays

For transient transformation of tobacco leaf epidermal cells, *BFP* or *LIR-SNT-BFP* and *GFP-ATG8a* were introduced into *pCAMBIA1302* vector under the *UBQ10* promoter. Soil-grown, 4-week-old *N. benthamiana* leaves were used for transient transformation. After two days of Agrobacterium inoculation, leaves were collected and cut into small squares for confocal microscopy. For transient transformation of Arabidopsis protoplasts, *BFP* or *LIR-SNT-BFP* were introduced into *pMD19* vector under the *UBQ10* promoter. Soil-grown, three-week-old transgenic Arabidopsis expressing *GFP-ATG8a* were used for preparing protoplasts. Transformation was done as described(Liu *et al*, 2022).

### Laser Scanning Confocal Microscopy (LSCM)

Lower epidermis of transformed *N.benthamiana* leaves, cotyledons of transgenic Arabidopsis seedlings and Arabidopsis protoplasts were observed with a Ni-E A1 HD25 confocal microscope (Nikon, Japan) and a STELLARIS 5 confocal microscope (Leica, Germany). Prior to image collection, the background auto-fluorescence was eliminated using untransformed samples. The BFP fluorescence signal was exited at 405 nm and emission was collected at 425-475 nm. The GFP fluorescence signal was exited at 488 nm and collected at 500-550 nm. The YFP fluorescence signal was exited at 514 nm and collected at 525-550 nm. The chlorophyll auto-fluorescence was exited with 640 nm laser and collected at 650-750 nm.

### Chloroplast isolation

Chloroplast isolation from Arabidopsis seedlings was performed as previously described(Wan *et al*., 2023). The chloroplasts were isolated from protoplasts through 40% and 85% Percoll step gradient, washed once with HEPES-sorbitol buffer, and processed for SDS-PAGE analysis.

### Immunoblotting and Co-immunoprecipitation

Protein extraction and immunoblotting were done as described(Luo *et al*., 2017). Semi-quantification of the protein levels was performed with ImageJ (https://imagej.nih.gov/) and protein levels were normalized to the H3. For immunoblotting, rabbit anti-H3 (1:8000 dilution, Abmart, China), mouse anti-GFP (1:5,000 dilution, Utibody, China), rabbit anti-tRFP (1:5000 dilution, Evrogen, Russia), mouse anti-RbcL (1:5000 dilution, Abmart, China), rabbit anti-RbcS (1:5000 dilution, Orizymes, China), rabbit anti-LHCA1 (1:5000 dilution, Orizymes, China), rabbit anti-Tic40 (1:2000 dilution, Agrisera, SWEDEN), rabbit anti-Toc75 (1:1000 dilution, gifted from Qihua Lin), rabbit anti-PDV2 (1:5000 dilution, gifted from Hongbo Gao), and the appropriate IgG-HRP conjugated secondary antibody (1:5000; ZSGB-Bio, China) were used. The signal was developed using Highly Sensitive ECL Chemiluminescence Substrate (LINDE, China) and chemiluminescence was detected using a chemiluminescent Western Blot scanner (ChemiScope 6100T, Clinx, China). All experiments were repeated at least three times with one representative result shown.

For co-immunoprecipitation, transgenic Arabidopsis co-expressing GFP-ATG8a and BFP-tagged proteins were used. Proteins were extracted with IP lysis buffer (50 mM Tris-HCl, pH 7.5, 150 mM NaCl, 1 mM MgCl_2_, 20% glycerol, 0.2% NP-40, and 1× protease inhibitor), and then centrifuged at 12000 rpm at 4°C for 10 min. The supernatant was incubated with GFP-beads for 3 h at 4°C with slow rotation. After three washes with washing buffer (50 mM Tris-HCl, pH 7.5, 150 mM NaCl, 1 mM MgCl_2_, 20% glycerol, 0.01% NP-40), bound proteins were eluted by boiling in SDS–PAGE loading buffer for immunoblotting.

### Measurement of the Number of Chloroplasts

First leaves of Arabidopsis seedlings were fixed in 3.5% (v/v) glutaraldehyde for 1 h in the dark and softened in 0.1 M Na_2_EDTA (pH 9) at 60°C for 3 h. DIC images were observed using a Ni-E A1 HD25 confocal microscope (Nikon, Japan).

### Chlorophyll measurements

Fresh leaves of Arabidopsis seedlings were collected, snap frozen and ground with liquid nitrogen. The powder was mixed with 80% acetone and incubated in the dark for 15-30 min. Cell debris was pelleted three times at 12,000 × g for 15 min at 4°C. The absorbance (A) of chlorophyll content was measured spectrophotometrically using 80% acetone as a blank control. The chlorophyll concentrations are calculated as follows: Chlorophyll *a* (mg/g) = [12.7 × A663 - 2.69 × A645] × V/W, Chlorophyll *b* (mg/g) = [22.9 × A645 - 4.86 × A663] × V/W, Chlorophyll *a+b* (mg/g) = [8.02 × A663 + 20.20 × A645] × V/W, Where V = volume of the extract (ml); W = weight of fresh leaves (mg).

### Transmission Electron Microscope (TEM)

Cotyledons of transgenic seedlings were fixed with 2.5% (v/v) glutaraldehyde overnight. Then samples were rinsed with 0.1 M phosphate buffer (Na_2_HPO_4_, NaH_2_PO_4_) and post-fixed with 1% OsO_4_ (w/v) for 2 h at 4°C. Following dehydration with alcohol and acetone series, samples were embedded in EPON 812 (Ted pella, USA) for 2 days at 60°C. Ultrathin sections (thickness 90 nm) were cut with a Leica EM UC7 (Leica, Germany), mounted on copper grids and contrasted with TI Blue stainer (Nisshin EM, Japan) and 3% lead citrate solution. The sections were visualized with a Tecnai G2 spirit Biotwin TEM (FEI, USA) at 120 kV accelerating voltage.

### Immunogold electron microscopy (IEM)

Cotyledons of transgenic seedlings were dissected and primary fixed in 0.25% glutaraldehyde and 1.5% paraformaldehyde in 50 mM PBS. Leaves were dehydrated in graded ethanol (20%, 40% for 30 min each) at 4℃ and secondary fixed in 0.5% osmium in 40% ethanol at -20℃ for 8 hours. Samples were further dehydrated in graded ethanol (75%, 95% and absolute ethanol for 45 min each) at 0℃ and infiltrated in LR white/ethanol graded mixture (1:3, 1:1, 3:1 and pure LR White for 45 min each) at room temperature. Samples were finally embedded in capsules with fresh LR white overnight at 60℃. The ultrathin sections (50 nm) were prepared and transferred to nickel grids. Double-immunogold labeling was performed ae previously described(Wang *et al*, 2016). The working concentration of primary antibodies (anti-GFP and anti-tRFP) were diluted 1:50. Gold particle coupled secondary anti-Mouse (10 nm, Sigma) and anti-Rabbit (15 nm, Sigma) antibodies were diluted 1:50, followed by the post-staining procedure by aqueous uranyl acetate/lead citrate. Ultrathin sections were examined using an FEI Talos L120C transmission electron microscope.

### Statistical analysis

Statistical analyses were performed using Student’s t-test. *, **, ***, ****, indicate significant difference with p < 0.05, p < 0.01, p < 0.001 and p < 0.0001, respectively. All data were presented as mean ± SE of at least three replicates, as indicated in the figure legends.

## Acknowledgements

We thank Cheng Chen (Shanghai Jiao Tong University) for *pdv2* seeds, Hongbo Gao (Beijing Forestry University) for PDV2 antibody, Qihua Ling (Institute of Plant Physiology and Ecology, Chinese Academy of Sciences) for anti-TOC75 antibody; Dr Yu Kong and Xu Wang (Electron Microscopy Facilities of Center for Excellence in Brain Science and Technology, Chinese Academy of Science) for assistance with Immuno-EM sample preparation and IEM image analysis. We thank Cheng Chen, Hongbo Gao, Qihua Ling, Quan-Sheng Qiu (Lanzhou University), Zhiping Xie (Shanghai Jiao Tong University), Heng Zhang (Shanghai Jiao Tong University), and Xin-Guang Zhu (Institute of Plant Physiology and Ecology, Chinese Academy of Sciences) for suggestions, and our lab members for discussions. This work was supported by grants from the National Natural Science Foundation of China (32270726, 92354302, 91954102, and 31871355), the Natural Science Foundation of Shanghai (23ZR1429400) to QG. TW is supported by Startup Fund for Young Faculty at SJTU (SFYF at SJTU).

## Author contributions

QG initiated the project. RL performed most of the experiments. XW, XL, YC, QL, LL, and DT performed some of the experiments. RL, TW, HW, ZK, and QG analyzed the data. RL and QG drafted the paper. QG edited the paper. TW and QG acquired funding. HW, ZK, and QG supervised the study.

## Declaration of interests

No conflict of interest is declared.

## Supplemental information titles and legends

**Figure S1 Protein sequences of LIR and SNT used to construct the synthetic chlorophagy receptor.**

**(A)** Schematic illustration of Arabidopsis NBR1 (uniprot ID: Q9SB64), with aa633-687 containing the LIR/AIM motif (WDPI, in red color) shown below. Residues W661 and I664 indicated with asterisks are required for interaction with ATG8. **(B)** Schematic illustration of AtSFR2 (uniprot ID: Q93Y07), with the N-terminal amphipathic helix (SNT), i.e. aa1-31, shown below. **(C)** Helical wheel projection of SNT. Clustering of hydrophobic residues in yellow color on one face of the helix suggests an amphipathic configuration.

**Figure S2 Transgenic lines carrying the synthetic chlorophagy receptor and a negative control in *GFP-ATG8a* background.**

**(A)** Arabidopsis transgenic lines (L1, L2, L16, L27, L32, and L36) expressing *LIR-SNT-BFP*, and transgenic lines (L1, L2, L6) expressing *SNT-BFP*, in the *GFP-ATG8a* background, were selected for further analysis. Five-day-old seedlings were treated with DMSO or 10 μM DEX for 10 days. Bar = 5 mm. **(B)** WB analysis of the protein expression levels in the transgenic lines in (A). Anti-GFP and Anti-tRFP were used to detect GFP-ATG8a, LIR-SNT-BFP and SNT-BFP. Histone H3 (H3) served as an internal control. **(C)** *LIR-SNT-BFP* was inserted in intergenic regions in L1, L2, and L27, as revealed by tail-PCR. The insertion sites and primers designed for genotyping are illustrated. **(D)** The T-DNA insertion sites of *GFP-ATG8a*, *GFP-ATG8a*/*LIR-SNT-BFP-*L1, L2, and L27 were verified by genomic PCR. Representative images or western blots from three biological replicates are shown in (A), (B), and (D).

**Fig S3 Induction of the synthetic chlorophagy receptor alone does not affect steady-state accumulation of chloroplast proteins.**

**(A)** Five-day-old seedlings expressing *LIR-SNT-BFP* in the *GFP-ATG8a* background, were treated with DMSO or 10 μM DEX for 10 days, before WB detection of chloroplast proteins TOC75 (chloroplast outer envelope protein), TIC40 (inner envelope protein), LHCA1 (thylakoid protein), RbcL (chloroplast-encoded stromal protein), and RbcS (stromal protein). Anti-GFP and Anti-tRFP were used to detect GFP-ATG8a and LIR-SNT-BFP. Histone H3 (H3) served as an internal control. Representative western blots from three biological replicates are shown. **(B)** Band intensities in (A) were quantified and normalized to H3. *GFP-ATG8a* in DMSO (DEX-) was set as 100%. Data are means ± SEM of three biological replicates. ns, no significant difference (student’s t-test). OE, chloroplast outer envelope; IE, chloroplast inner envelope; Thy, thylakoids; St, stroma.

**Fig S4 SNT-BFP-LIR facilitates chloroplast autophagic degradation similar as LIR-SNT-BFP.**

**(A)** Co-immunoprecipitation assay showing the *in vivo* interaction between GFP-ATG8a and SNT-BFP-LIR. Transgenic lines co-expressing *GFP-ATG8a* and *BFP*, *SNT-BFP*, *LIR-SNT-BFP*, or *SNT-BFP-LIR* were treated with 30 μM DEX before co-IP. Proteins were immunoprecipitated with GFP-Trap beads and detected with anti-GFP and anti-tRFP. **(B)** Five-day-old transgenic lines (L1, L13, L21) expressing *SNT-BFP-LIR* in the *GFP-ATG8a* background, were treated with DMSO or 10 μM DEX for 10 days. Bar = 5 mm. **(C)** SNT-BFP-LIR promotes the degradation of chloroplast proteins in the presence of Lin. Five-day-old seedlings expressing *SNT-BFP-LIR* in the *GFP-ATG8a* background, were treated with DMSO or 10 μM DEX in the presence of 0.5 mM Lin for 14 days, before WB detection of chloroplast proteins TOC75 (chloroplast outer envelope protein), TIC40 (inner envelope protein), LHCA1 (thylakoid protein), RbcL (chloroplast encoded stromal protein), and RbcS (stromal protein). Anti-GFP and Anti-tRFP were used to detect GFP-ATG8a and LIR-SNT-BFP. Histone H3 (H3) served as an internal control. **(D)** Band intensities in (C) were quantified and normalized to H3. *GFP-ATG8a* in DMSO (DEX-) was set as 100%. Data are means ± SEM of three biological replicates. **, *p* < 0.01; *, *p* < 0.05; ns, no significant difference (student’s t-test). Representative images or western blots from three biological replicates are shown in (A), (B), and (C). OE, chloroplast outer envelope; IE, chloroplast inner envelope; Thy, thylakoids; St, stroma.

**Fig S5 Response of DEX-treated *GFP-ATG8a* and *GFP-ATG8a/LIR-SNT-BFP* to herbicides.**

**(A)** Both NF and DCMU triggered cell death in the cotyledon mesophyll cells, which was visualized by propidium iodine (PI) staining. Five-day-old *GFP-ATG8a* were treated with 10 μM DEX, or 1 μM NF, or 2 μM DCMU, or combinations of the chemicals for 10 days, before confocal imaging. Bar = 10 μm. **(B)** Both NF and DCMU triggered abnormal and uneven distribution of chlorophyll. Five-day-old *GFP-ATG8a* were treated with 10 μM DEX, or 1 μM NF, or 10 μM DCMU, or combinations of the chemicals for 3 days, before confocal imaging. Chlorophyll fluorescence along the white lines plotted to the right. Bar = 5 μm. **(C)** Seedlings grown in soil were partially suppressed of NF-induced plant death by LIR-SNT-BFP induction. Transgenic seeds carrying *GFP-ATG8a/LIR-SNT-BFP* were germinated in soil, and started treatments at growth stage 1.02 (day 8). Seedlings in some pots were sprayed with 1 μM NF daily for 3 or 5 days, meanwhile they were also sprayed with 10 μM DEX daily for 7 days. All seedlings were photographed at day 15. Bar = 1 cm. Representative images from three biological replicates are shown in (A), (B), and (C).

**Fig S6 The synthetic chlorophagy receptor partially protects *pdv2* from norflurazon.**

**(A)** LIR-SNT-BFP induction partially protected cotyledons from chlorosis in NF-treated *pdv2*. Five-day-old *pdv2*, *GFP-ATG8a/LIR-SNT-BFP*, and *pdv2/GFP-ATG8a/LIR-SNT-BFP* were treated with 10 μM DEX, or 1 μM NF, or DEX plus NF for 10 days. Bar = 2 mm. **(B)** LIR-SNT-BFP induction partially protected chlorophyll loss in NF-treated *pdv2*. Cotyledon mesophyll cells from plants in (A) were analyzed by confocal microscopy. Bar = 10 μm. Representative images from three biological replicates are shown in (A) and (B).

**Fig S7 Chloroplast sizes were reduced in *atg5* and *atg7* upon induced expression of the synthetic chlorophagy receptor.**

**(A)** PCR-verification of atg2/GFP-ATG8a and atg2/GFP-ATG8a/LIR-SNT-BFP. **(B)** The expression of LIR-SNT-BFP is silenced at the protein level in the *atg2* background. WB analysis of five-day-old seedlings expressing *GFP-ATG8a* or *GFP-ATG8a/LIR-SNT-BFP* in the *atg2* background, treated with DMSO or 10 μM DEX for 10 days. Anti-GFP and Anti-tRFP were used to detect GFP-ATG8a and LIR-SNT-BFP. Histone H3 (H3) served as an internal control. **(C)** PCR-verification of *atg5/GFP-ATG8a*, *atg5/GFP-ATG8a/LIR-SNT-BFP*, *atg7/GFP-ATG8a* and *atg7/GFP-ATG8a/LIR-SNT-BFP*. **(D)** LIR-SNT-BFP induction reduces the sizes of chloroplasts in *atg5* and *atg7*. Cotyledons of five-day-old *GFP-ATG8a*, *GFP-ATG8a/LIR-SNT-BFP*, *atg5/GFP-ATG8a/LIR-SNT-BFP* and *atg7/GFP-ATG8a/LIR-SNT-BFP*, treated with DMSO or 10 μM DEX for 6 days, were analyzed by TEM. Bar = 1 μm. **(E)** Quantification of the lengths of chloroplasts in (D). Data are means ± SEM (n = 40). ****, *p* < 0.0001; ns, no significant difference (student’s t-test). Representative images or western blots from three biological replicates are shown in (A), (B), and (C).

**Table S1. Primers used in this study.**

**Movies1-4.**

**Movie 1 YFP-VAMP711-labelled tonoplast quietly surrounds chloroplasts in DMSO-treated *GFP-ATG8a/LIR-SNT-BFP* leaf mesophyll cells.**

**Movie 2 YFP-VAMP711-labelled tonoplast quietly surrounds chloroplasts in DMSO-treated *GFP-ATG8a/LIR-SNT-BFP* leaf mesophyll cells (a second area).**

**Movie 3 Dynamic interactions between YFP-VAMP711-labelled tonoplast and chloroplasts in DEX-treated *GFP-ATG8a/LIR-SNT-BFP* leaf mesophyll cells.**

**Movie 4 Dynamic vacuole fission and fusion in DEX-treated *GFP-ATG8a/LIR-SNT-BFP* leaf mesophyll cells.**

## References

Aoyama T, Chua NH (1997) A glucocorticoid-mediated transcriptional induction system in transgenic plants. Plant J 11: 605–612

Carrión CA, Costa ML, Martínez DE, Mohr C, Humbeck K, Guiamet JJ (2013) inhibition of cysteine proteases provides evidence for the involvement of ’senescence-associated vacuoles’ in chloroplast protein degradation during dark-induced senescence of tobacco leaves. J Exp Bot 64: 4967–4980

Chang N, Sun QQ, Li YQ, Mu YJ, Hu JL, Feng Y, Liu XM, Gao HB (2017) PDV2 has a dosage effect on chloroplast division in. Plant Cell Rep 36: 471–480

Chen G, Kroemer G, Kepp O (2020) Mitophagy: An Emerging Role in Aging and Age-Associated Diseases. Front Cell Dev Biol 8

Fourrier N, Bédard J, Lopez-Juez E, Barbrook A, Bowyer J, Jarvis P, Warren G, Thorlby G (2008) A role for SENSITIVE TO FREEZING2 in protecting chloroplasts against freeze-induced damage in Arabidopsis. Plant J 55: 734–745

Frank M, Duvezin-Caubet S, Koob S, Occhipinti A, Jagasia R, Petcherski A, Ruonala MO, Priault M, Salin B, Reichert AS (2012) Mitophagy is triggered by mild oxidative stress in a mitochondrial fission dependent manner. Bba-Mol Cell Res 1823: 2297–2310

Fukuda T, Furukawa K, Maruyama T, Yamashita SI, Noshiro D, Song C, Ogasawara Y, Okuyama K, Alam JM, Hayatsu M et al (2023) The mitochondrial intermembrane space protein mitofissin drives mitochondrial fission required for mitophagy. Mol Cell 83: 2045–2058.e2049

Geldner N, Dénervaud-Tendon V, Hyman DL, Mayer U, Stierhof YD, Chory J (2009) Rapid, combinatorial analysis of membrane compartments in intact plants with a multicolor marker set. Plant J 59: 169–178

Gubas A, Dikic I (2022) A Guide To The regulation of selective autophagy receptors. Febs J 289: 75–89

Ishida H, Yoshimoto K, Izumi M, Reisen D, Yano Y, Makino A, Ohsumi Y, Hanson MR, Mae T (2008) Mobilization of rubisco and stroma-localized fluorescent proteins of chloroplasts to the vacuole by an ATG gene-dependent autophagic process. Plant Physiol 148: 142–155

Izumi M, Ishida H, Nakamura S, Hidema J (2017) Entire Photodamaged Chloroplasts Are Transported to the Central Vacuole by Autophagy. Plant Cell 29: 377–394

Izumi M, Nakamura S (2018) Chloroplast Protein Turnover: The Influence of Extraplastidic Processes, Including Autophagy. Int J Mol Sci 19

Izumi M, Nakamura S, Otomo K, Ishida H, Hidema J, Nemoto T, Hagihara S (2024) Autophagosome development and chloroplast segmentation occur synchronously for piecemeal degradation of chloroplasts. Elife 12

Ji CH, Kim HY, Lee MJ, Heo AJ, Park DY, Lim S, Shin S, Ganipisetti S, Yang WS, Jung CA et al (2022) The AUTOTAC chemical biology platform for targeted protein degradation via the autophagy-lysosome system (vol 13, 904, 2022). Nat Commun 13

Kikuchi Y, Nakamura S, Woodson JD, Ishida H, Ling Q, Hidema J, Jarvis RP, Hagihara S, Izumi M (2020) Chloroplast Autophagy and Ubiquitination Combine to Manage Oxidative Damage and Starvation Responses. Plant Physiol 183: 1531–1544

Klionsky DJ, Petroni G, Amaravadi RK, Baehrecke EH, Ballabio A, Boya P, Bravo-San Pedro JM, Cadwell K, Cecconi F, Choi AMK et al (2021) Autophagy in major human diseases. Embo J 40

Lee HN, Chacko JV, Solis AG, Chen KE, Barros JA, Signorelli S, Millar AH, Vierstra RD, Eliceiri KW, Otegui MS et al (2023) The autophagy receptor NBR1 directs the clearance of photodamaged chloroplasts. Elife 12

Lemke MD, Fisher KE, Kozlowska MA, Tano DW, Woodson JD (2021) The core autophagy machinery is not required for chloroplast singlet oxygen-mediated cell death in the Arabidopsis thaliana plastid ferrochelatase two mutant. BMC Plant Biol 21: 342

Li Z, Wang C, Wang Z, Zhu C, Li J, Sha T, Ma L, Gao C, Yang Y, Sun Y et al (2019) Allele-selective lowering of mutant HTT protein by HTT-LC3 linker compounds. Nature 575: 203–209

Linnane AW, Stewart PR (1967) The inhibition of chlorophyll formation in Euglena by antibiotics which inhibit bacterial and mitochondrial protein synthesis. Biochem Biophys Res Commun 27: 511–516

Liu M, Yu J, Yang M, Cao L, Chen C (2024) Adaptive evolution of chloroplast division mechanisms during plant terrestrialization. Cell Rep 43: 113950

Liu R, Zhang R, Yang Y, Liu X, Gong Q (2022) Monitoring Autophagy in Rice With GFP-ATG8 Marker Lines. Frontiers in Plant Science 13

Luo L, Zhang P, Zhu R, Fu J, Su J, Zheng J, Wang Z, Wang D, Gong Q (2017) Autophagy Is Rapidly Induced by Salt Stress and Is Required for Salt Tolerance in Arabidopsis. Front Plant Sci 8: 1459

Luo MQ, Law KC, He YL, Chung KK, Po MK, Feng LL, Chung KP, Gao CJ, Zhuang XH, Jiang LW (2023a) Arabidopsis AUTOPHAGY-RELATED2 is essential for ATG18a and ATG9 trafficking during autophagosome closure. Plant Physiol

Luo N, Shang DD, Tang ZW, Mai JY, Huang X, Tao LZ, Liu LC, Gao CJ, Qian YW, Xie QJ et al (2023b) Engineered ATG8-binding motif-based selective autophagy to degrade proteins and organelles. New Phytol 237: 684–697

Martinez DE, Costa ML, Gomez FM, Otegui MS, Guiamet JJ (2008) ’Senescence-associated vacuoles’ are involved in the degradation of chloroplast proteins in tobacco leaves. Plant J 56: 196–206

Mei LG, Chen XR, Wei FJ, Huang X, Liu L, Yao J, Chen J, Luo XG, Wang ZL, Yang AM (2023) Tethering ATG16L1 or LC3 induces targeted autophagic degradation of protein aggregates and mitochondria. Autophagy 19: 2997–3013

Michaeli S, Honig A, Levanony H, Peled-Zehavi H, Galili G (2014) ATG8-INTERACTING PROTEIN1 Is Involved in Autophagy-Dependent Vesicular Trafficking of Plastid Proteins to the Vacuole. Plant Cell 26: 4084–4101

Miyagishima SY, Froehlich JE, Osteryoung KW (2006) PDV1 and PDV2 mediate recruitment of the dynamin-related protein ARC5 to the plastid division site. Plant Cell 18: 2517–2530

Nakamura S, Hidema J, Sakamoto W, Ishida H, Izumi M (2018) Selective Elimination of Membrane-Damaged Chloroplasts via Microautophagy. Plant Physiol 177: 1007–1026

Noda NN, Ohsumi Y, Inagaki F (2010) Atg8-family interacting motif crucial for selective autophagy. FEBS Lett 584: 1379–1385

Okazaki K, Kabeya Y, Suzuki K, Mori T, Ichikawa T, Matsui M, Nakanishi H, Miyagishima S (2009a) The PLASTID DIVISION1 and 2 Components of the Chloroplast Division Machinery Determine the Rate of Chloroplast Division in Land Plant Cell Differentiation. Plant Cell 21: 1769–1780

Okazaki K, Kabeya Y, Suzuki K, Mori T, Ichikawa T, Matsui M, Nakanishi H, Miyagishima SY (2009b) The PLASTID DIVISION1 and 2 components of the chloroplast division machinery determine the rate of chloroplast division in land plant cell differentiation. Plant Cell 21: 1769–1780

Ono Y, Wada S, Izumi M, Makino A, Ishida H (2013) Evidence for contribution of autophagy to rubisco degradation during leaf senescence in Arabidopsis thaliana. Plant Cell Environ 36: 1147–1159

Osteryoung KW, Stokes KD, Rutherford SM, Percival AL, Lee WY (1998) Chloroplast division in higher plants requires members of two functionally divergent gene families with homology to bacterial ftsZ. Plant Cell 10: 1991–2004

Otegui MS, Noh YS, Martínez DE, Vila Petroff MG, Andrew Staehelin L, Amasino RM, Guiamet JJ (2005) Senescence-associated vacuoles with intense proteolytic activity develop in leaves of Arabidopsis and soybean. Plant J 41: 831–844

Pan T, Liu Y, Hu X, Li P, Lin C, Tang Y, Tang W, Liu Y, Guo L, Kim C et al (2023) Stress-induced endocytosis from chloroplast inner envelope membrane is mediated by CHLOROPLAST VESICULATION but inhibited by GAPC. Cell Rep 42: 113208

Park JH, Tran LH, Jung S (2017) Perturbations in the Photosynthetic Pigment Status Result in Photooxidation-Induced Crosstalk between Carotenoid and Porphyrin Biosynthetic Pathways. Front Plant Sci 8

Raffeiner M, Zhu SS, González-Fuente M, Üstün S (2023) Interplay between autophagy and proteasome during protein turnover. Trends Plant Sci 28: 698–714

Ridley SM (1977) Interaction of Chloroplasts with Inhibitors - Induction of Chlorosis by Diuron during Prolonged Illumination Invitro. Plant Physiol 59: 724–732

Rogov VV, Nezis IP, Tsapras P, Zhang H, Dagdas Y, Noda NN, Nakatogawa H, Wirth M, Mouilleron S, McEwan DG et al (2023) Atg8 family proteins, LIR/AIM motifs and other interaction modes. Autophagy Rep 2

Sakai Y, Koller A, Rangell LK, Keller GA, Subramani S (1998) Peroxisome degradation by microautophagy in Pichia pastoris: identification of specific steps and morphological intermediates. J Cell Biol 141: 625–636

Svenning S, Lamark T, Krause K, Johansen T (2011) Plant NBR1 is a selective autophagy substrate and a functional hybrid of the mammalian autophagic adapters NBR1 and p62/SQSTM1. Autophagy 7: 993–1010

Takahashi D, Moriyama J, Nakamura T, Miki E, Takahashi E, Sato A, Akaike T, Itto-Nakama K, Arimoto H (2019) AUTACs: Cargo-Specific Degraders Using Selective Autophagy. Mol Cell 76: 797–810.e710

Tan SX, Wang D, Fu YH, Zheng HW, Liu Y, Lu BX (2023) Targeted clearance of mitochondria by an autophagy-tethering compound (ATTEC) and its potential therapeutic effects. Sci Bull 68: 3013–3026

Thompson AR, Doelling JH, Suttangkakul A, Vierstra RD (2005) Autophagic nutrient recycling in Arabidopsis directed by the ATG8 and ATG12 conjugation pathways. Plant Physiol 138: 2097–2110

Wada S, Ishida H, Izumi M, Yoshimoto K, Ohsumi Y, Mae T, Makino A (2009) Autophagy plays a role in chloroplast degradation during senescence in individually darkened leaves. Plant Physiol 149: 885–893

Wan C, Zhang H, Cheng H, Sowden RG, Cai W, Jarvis RP, Ling Q (2023) Selective autophagy regulates chloroplast protein import and promotes plant stress tolerance. Embo j 42: e112534

Wang H, Zhuang X, Wang X, Law AHY, Zhao T, Du S, Loy MMT, Jiang L (2016) A Distinct Pathway for Polar Exocytosis in Plant Cell Wall Formation. Plant Physiology 172: 1003–1018

Wang S, Blumwald E (2014) Stress-induced chloroplast degradation in Arabidopsis is regulated via a process independent of autophagy and senescence-associated vacuoles. Plant Cell 26: 4875–4888

Wang Y, Yu BJ, Zhao JP, Guo JB, Li Y, Han SJ, Huang L, Du YM, Hong YG, Tang DZ et al (2013) Autophagy Contributes to Leaf Starch Degradation. Plant Cell 25: 1383–1399

Yamamoto H, Zhang SD, Mizushima N (2023) Autophagy genes in biology and disease. Nat Rev Genet 24: 382–400

Yan H, Qi A, Lu Z, You Z, Wang Z, Tang H, Li X, Xu Q, Weng X, Du X et al (2024) Dual roles of AtNBR1 in regulating selective autophagy via liquid-liquid phase separation and recognition of non-ubiquitinated substrates in Arabidopsis. Autophagy 20: 2804–2815

Zhuang XH, Jiang LW (2019) Chloroplast Degradation: Multiple Routes Into the Vacuole. Front Plant Sci 10

Žnidarič MT, Zagorščak M, Ramšak Ž, Stare K, Chersicola M, Novak MP, Kladnik A, Dermastia M (2025) Chloroplast Vesiculation and Induced Chloroplast Vesiculation and Senescence-Associated Gene 12 Expression During Tomato Flower Pedicel Abscission. Plant Direct 9: e70035

